# Nazo, the *Drosophila* homolog of the NBIA-mutated protein – c19orf12, is required for triglyceride homeostasis

**DOI:** 10.1101/2022.09.29.510106

**Authors:** Perinthottathil Sreejith, Sara Lolo, Kristen R. Patten, Maduka Gunasinghe, Neya More, Leo J. Pallanck, Rajnish Bharadwaj

**Affiliations:** Department of Pathology and Laboratory Medicine, University of Rochester Medical Center, Rochester, NY, USA; Department of Genome Sciences, University of Washington, Seattle, WA, USA

**Keywords:** c19orf12, NBIA, Nazo, CG3740, lipid homeostasis, neurodegeneration, membrane contact site, ER-lipid droplet contact site, lipid droplet

## Abstract

Lipid dyshomeostasis has been implicated in a variety of diseases ranging from obesity to neurodegenerative disorders such as NBIA. Here, we uncover the physiological role of Nazo, the *Drosophila* homolog of the NBIA-mutated protein – c19orf12, whose function has been elusive. Ablation of *Drosophila* c19orf12 homologs leads to dysregulation of multiple lipid metabolism genes. *nazo* mutants exhibit markedly reduced gut lipid droplet and whole-body triglyceride contents. Consequently, they are sensitive to starvation and oxidative stress. Nazo localizes to ER-lipid droplet contact sites and is required for maintaining normal levels of Perilipin2, an inhibitor of the lipase – Brummer. Concurrent knockdown of Brummer or overexpression of Perilipin2 rescues the *nazo* phenotype, suggesting that this defect may arise from diminished Perilipin2 on lipid droplets leading to aberrant Brummer-mediated lipolysis. Our findings provide novel insights into the role of c19orf12 as a possible link between lipid dyshomeostasis and neurodegeneration, particularly in the context of NBIA.

## INTRODUCTION

Triglycerides and their numerous lipid derivatives (e.g., fatty acids, phospholipids) are fundamental to life as they serve diverse biological functions such as providing a source of energy, acting as building blocks for cell membranes and modulating signaling pathways ^1,2^. Therefore, proper intake, synthesis, storage, transport, and metabolism of triglycerides are highly regulated processes that have profound impacts on animal health. Defects in triglyceride hemostasis have been implicated in numerous diseases including obesity, diabetes, and neurodegenerative disorders ^1,3-5^.

Triglyceride metabolic pathways are highly conserved throughout evolution, and animal models such as *Drosophila melanogaster* have been instrumental in advancing our understanding of lipid homeostasis ^2,6^. Like human liver and intestine, *Drosophila* gut serves as a crucial hub for triglyceride homeostasis by orchestrating its absorption, storage, and distribution to other organs. Triglycerides derived from the food are converted into fatty acids in the gut by intestinal lipases. Fatty acids are absorbed by the enterocytes, which reconvert them into triglycerides and store them in intestinal lipid droplets ^2^. In addition, de novo synthesis of fatty acids and triglycerides from glucose serves as an alternative source of gut triglycerides. For mobilization out of enterocytes, triglycerides are hydrolyzed into diacylglycerol, which are subsequently packaged into lipoprotein particles and transported to fat body and other tissues via hemolymph ^2^.

The biochemical pathway for triacylglycerol synthesis from fatty acids is well characterized, and involves the enzymes GPAT, AGPAT, PAP and DGAT, which are conserved in *Drosophila* ^2^. Triacylglycerol (TAG) synthesis occurs between endoplasmic reticulum (ER) leaflets at discrete subdomains delineated by proteins like Seipin ^7-10^. Subsequently, triglycerides are packaged into lipid droplets (LD), which bud off from the endoplasmic reticulum but maintain physical association with ER through endoplasmic reticulum-lipid droplet (ER-LD) contact sites ^10^. Like membrane contact sites between other organelles (e.g., ER and mitochondria), ER-LD contact sites serve as a direct conduit for transfer of lipids and proteins between the organelles they bridge. Their disruption results in dramatic lipid droplet abnormalities both at the morphological and proteome level ^11,12^. Their significance is underscored by the fact that, mutations in genes encoding ER-LD contact site proteins like Seipin lead to a wide variety of deficits ranging from lipodystrophy to neurodegeneration ^3,4^.

Depending on cellular and organismal demands, triglycerides are converted into fatty acids either by cytosolic lipases (e.g., ATGL/brummer) or lipophagy ^13,14^. Fatty acids in turn can be channeled into various pathways such as Beta oxidation for energy production or Kennedy pathway for synthesis of phospholipids ^15^. In *Drosophila* the lipase Brummer plays a key role in lipolysis, and not surprisingly, *brummer* mutants are characterized by aberrant triglyc ride accumulation ^13^. A critical regulator of Brummer activity is the lipid droplet protein Perilipin 2, which maintains normal triglyceride stores by antagonizing Brummer ^16^.

Both catabolic and anabolic steps in triglycerides metabolism are exquisitely regulated by a battery of hormones such as insulin like peptides (DILPs) and adipokinetic hormone ^1,17,18^. In addition, gut microbiota influence triglyceride homeostasis through modulation of various signaling pathways including the innate immune response NF-κB pathway and insulin pathway ^19-21^. Interestingly, recent evidence suggests that the Drosophila NF-κB homolog Relish and its downstream effector – Sting can affect fat body triglyceride levels, independent of their roles in immune response ^22,23^.

The focus of the current study is the Drosophila homologs of human *c19orf12* gene, which is mutated in a familial neurodegenerative disorder – Neurodegeneration with brain iron accumulation (NBIA) ^24-26^. As the name of the disease suggests, NBIA patients show prominent neuronal loss accompanied by iron accumulation in basal ganglia. However, these patients also suffer from neuronal dysfunction/loss involving other areas of brain that do not exhibit significant iron accumulation, suggesting that the primary role of c19orf12 may be outside iron homeostasis. In line with this, other NBIA-mutated genes participate in diverse biological pathways including lipid metabolism, mitochondrial function, autophagolysosomal degradation and iron metabolism ^24,26^.

The biological role of c19orf12 is largely unknown. Human c19orf12 is a widely expressed gene, which is present in brain but strikingly enriched in adipose tissue (based on human protein atlas, Supplementary figure. 3). In cell culture, it localizes to endoplasmic reticulum and mitochondria ^27^. In concordance with this, patient derived fibroblasts from c19orf12 patients show increased sensitivity to oxidative stress and increased mitochondrial calcium in response to ATP stimulation ^27^. In addition, a recent study has demonstrated that c19orf12 knockout in M17 neuronal cells causes cytosolic iron accumulation and increased sensitivity to ferroptosis ^28^. The only studies on this gene on an organismal level has been performed in *Drosophila melanogaster*. c19orf12 is evolutionarily conserved and *Drosophila melanogaster* has two homologs of this gene named – CG3740 and Nazo (CG11671). Using RNAi transgenic lines, Iuso et al showed that depletion of fly c19orf12 homologs results in neurodegeneration and bang sensitivity ^29^, a commonly used phenotype in Drosophila to assess neuronal function ^29^. In an unrelated context, Goto et al demonstrated that the fly c19orf12 homolog – Nazo is involved in NFkB-mediated antiviral response ^30^.

In this paper, we identify a novel function for the Drosophila c19orf12 homolog – Nazo in lipid homesotasis. We demonstrate that nazo mutants exhibit extensive lipid droplet depletion in gut. Consequently, these mutants have a diminished lifespan and are sensitive to starvation. Nazo is an ER-lipid droplet contact site protein, which inhibits lipolysis by keeping the lipase activity of Brummer in check. These findings are particularly interesting considering the accumulating evidence linking lipid dyshomeostasis to NBIA and other forms of neurodegeneration. Our study provides novel insights into the physiological function of c19orf12 and the molecular basis of NBIA.

## RESULTS

### *Drosophila nazo* mutants show diminished longevity and functional deficits

As previously described, human c19orf12 gene has two *Drosophila melanogaster* orthologs – CG3740 and Nazo ^29^. To investigate the role of *Drosophila* c19orf12 orthologs, we generated deletion mutants for these genes using CRISPR-CAS9 based genome engineering approach.

The *nazo* mutant has a 657-nucleotide deletion that includes the translation initiation site and majority of the open reading frame. The *CG3740* mutant allele consists of a 134-nucleotide deletion, which most likely represents a null mutant as it includes the transmembrane domain and introduces a frameshift mutation into the downstream open reading frame (Fig. 1A). Homozygous mutant flies for both Nazo and CG3740 are viable. However, *nazo* as well as nazo/*CG3740* double mutants demonstrate a marked reduction in longevity with a median life span of about 40 days as compared to 60 days for the wild type flies (Fig 1B). The deletion of CG3740 has subtle effects on longevity (Fig. 1B). To assess the functional significance of these mutations, we examined the locomotor abilities of these mutants using a standard climbing assay. At the age of 20 days, *nazo* single mutants as well as *CG3740/nazo* double mutants exhibit a markedly compromised locomotor ability (Fig. 1C). The average distance climbed during the assay is 267.5 ± 28.4 mm for 20-day-old WT flies as opposed to 99.5 ± 18.6 mm and 79.7± 7.3 mm for *nazo* and *nazo/CG3740* double mutants, respectively (Fig. 1C). Overall, these data demonstrate that the Drosophila c19orf12 ortholog Nazo is required for normal lifespan and fitness of the animal.

**Figure 1.**
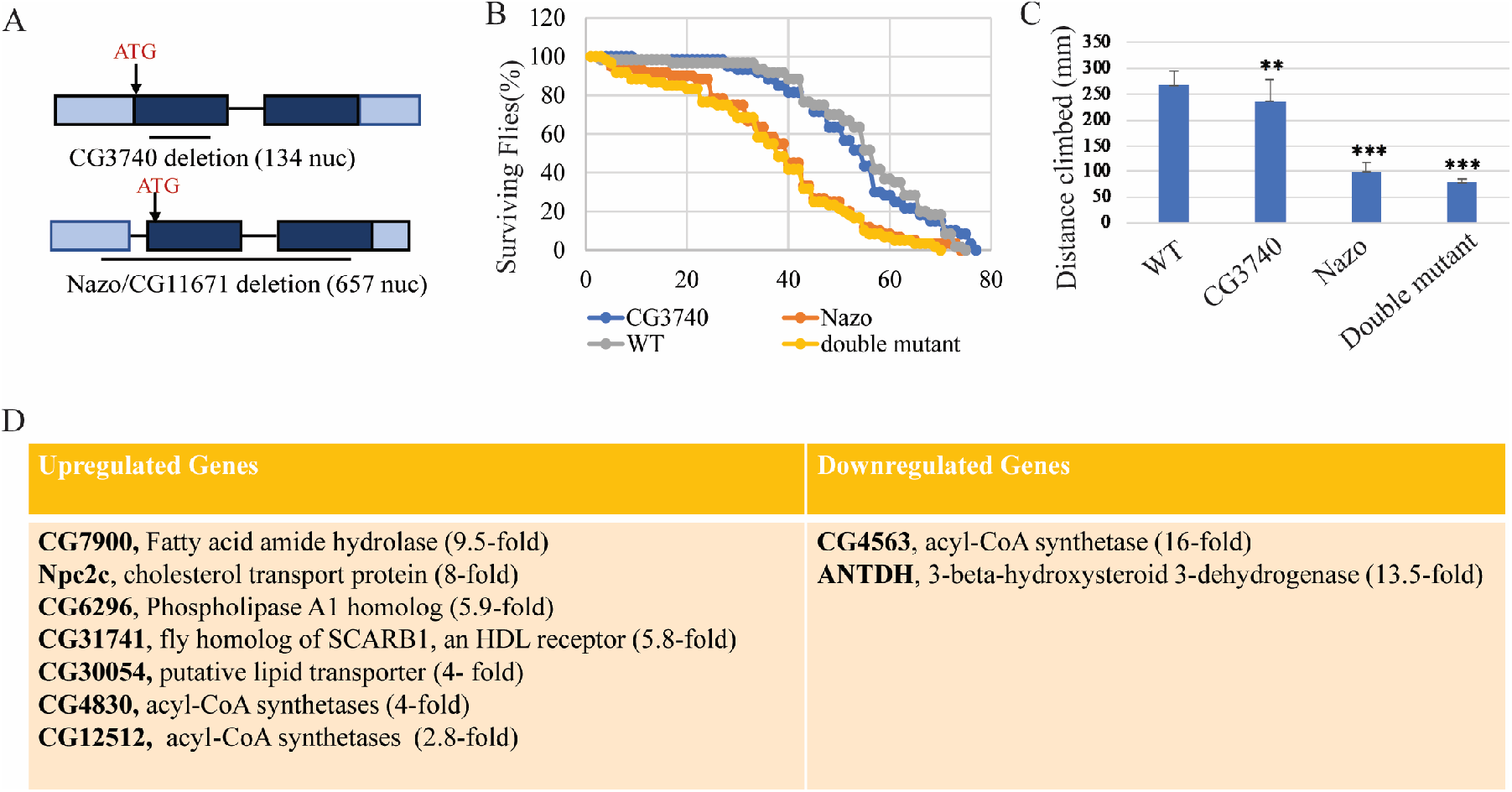
Drosophila C19orf12 mutants show reduced lifespan, functional deficits, and dysregulation of lipid metabolism genes. (A) Schematic of *Drosophila* Nazo and CG3740 genomic locus with deletion (open reading frame in dark blue, 5” and 3” untranslated regions in light blue, extent of deletion indicated by line) (B) Lifespan of males of indicated genotypes (N=60) (C) Standard climbing assay performed on 20-day old flies of indicated genotypes. Data are mean ± s.e.m. (*N* = 60 for each genotype, \*\*\**p* < 0.005 by Student’s *t*-test). (D) Lipid metabolism genes upregulated or downregulated (with indicated fold change) in guts of 20-day old *nazo/CG3740* mutant males relative to WT flies

### *nazo* mutants exhibit gut lipid droplet depletion

According to modeENCODE tissue expression data, while CG3740 is variably expressed in multiple tissues (including brain and gut), Nazo is predominantly expressed in gut. To gain insight into the physiological role of Nazo, we performed RNA seq analysis on wild type and *nazo/CG3740* double mutant male guts (where both these proteins are highly enriched). We identified 248 genes that were upregulated and 184 genes that were downregulated more than two folds in guts from nazo/*CG3740* double mutants relative to WT 20-day-old flies (Supplemental table 1 and 2). Interestingly, many of the top hits from this analysis were genes involved in lipid metabolism (Fig. 1D). Other cell biological processes significantly represented by the differentially expressed genes include immune response, metal homeostasis, endoplasmic reticulum stress and transcriptional regulation.

Prompted by this finding, we examined lipid droplets in Drosophila guts by BODIPY™ 493/503 dye staining. This revealed extensive lipid depletion in guts from nazo/CG3740 double mutant males relative to WT males (20-day-old) with 100% penetrance (Fig. 2A, B). The distribution of lipid droplets in female guts is different from the male guts; in male guts lipid droplets are predominantly confined to the R4 and R5 regions, whereas in females their distribution is more diffuse. The lipid droplet depletion is also present in mutant females but less pronounced than males. The severity and penetrance of this phenotype is similar in *nazo* single and *nazo/CG3740* double mutants suggesting that Nazo plays the primary role in this context. We also assessed the impact of starvation on this phenotype. We examined the gut lipid content of 20-day-old WT and mutant males, after 40 hours of starvation, using BODIPY™ 493/503 staining. The gut lipid droplet depletion was even more dramatic in the *nazo* mutants and double mutants under starved condition (Fig. 2C). Overall, our findings indicate that Nazo is required for the maintenance of lipid droplets in gut.

**Figure 2.**
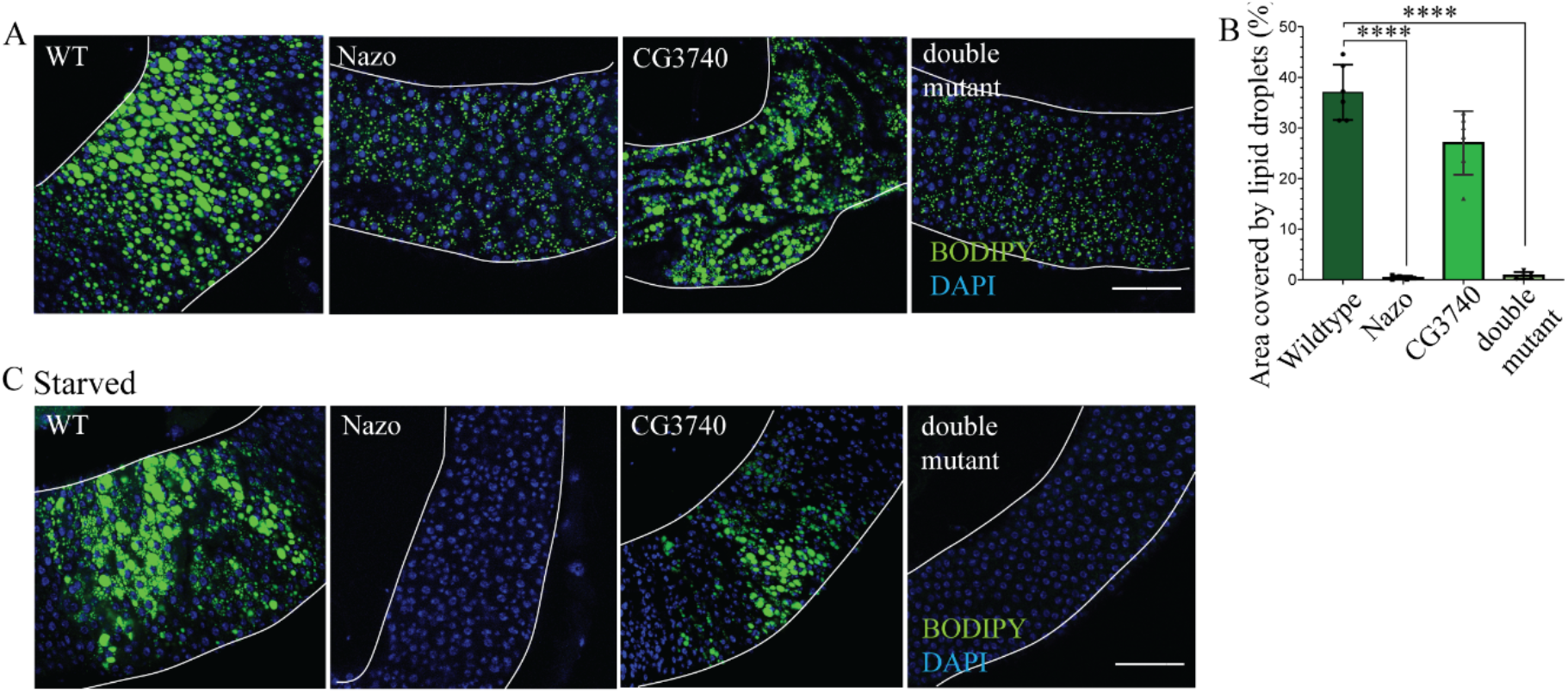
*nazo* mutants show gut lipid depletion. (A) Confocal images of guts from 20-day old flies stained with BODIPY™ 493/503 dye show markedly reduced lipid droplet content in midguts. (BODIPY™ 493/503 dye – green and DAPI – blue; Scale bar = 10µm) (B) Quantification of percentage area occupied by lipid droplets in the midgut R4 region (N=20; Student’s t-test **** p-value =< 0.0001) (C) Guts from 20-day old flies stained with BODIPY™ 493/503 dye after 40 hours of starvation show almost complete loss of lipid droplets in *nazo* and *nazo/CG3740* double mutants. (BODIPY™ 493/503 dye – green and DAPI – blue; Scale bar = 10µm)

### Nazo gut lipid depletion phenotype is not due to disrupted immune response

Multiple studies have shown that bacterial infection influences gut and fat body lipid droplet levels ^21^. Furthermore, Nazo is induced by the fly NF-kB pathway and plays a role in viral immune response in S2 cells ^30^. Though the lipid depletion phenotype we observed was in the absence of any exogenously introduced bacterial infection, we asked if this phenotype was related to impaired immune response. To this end, we performed lipid droplet staining on axenic (germ-free) WT and mutant flies generated through antibiotic treatment ^31^.

We found that under axenic conditions the *nazo* and *nazo/CG3740* double mutant flies showed similar lipid droplet depletion phenotypes as seen in untreated condition (Fig. 3A, B). Given the above-mentioned connection between NF-kB pathway and nazo, and the fat body lipid depletion phenotype observed in *relish* ^23^ and *sting* ^22^ mutants, we also performed lipid droplet staining on guts from flies lacking these proteins. Gut lipid droplet depletion was not observed in *relish* mutants or in flies with actin-Gal4 driven ubiquitous expression of sting RNAi (data not shown). The latter observation was consistent with what was seen for *sting* mutant by Akhmetova *et al* ^22^. Collectively, these findings suggest that the lipid depletion phenotype observed in *nazo* mutants is not secondary to compromised immune response.

**Figure 3.**
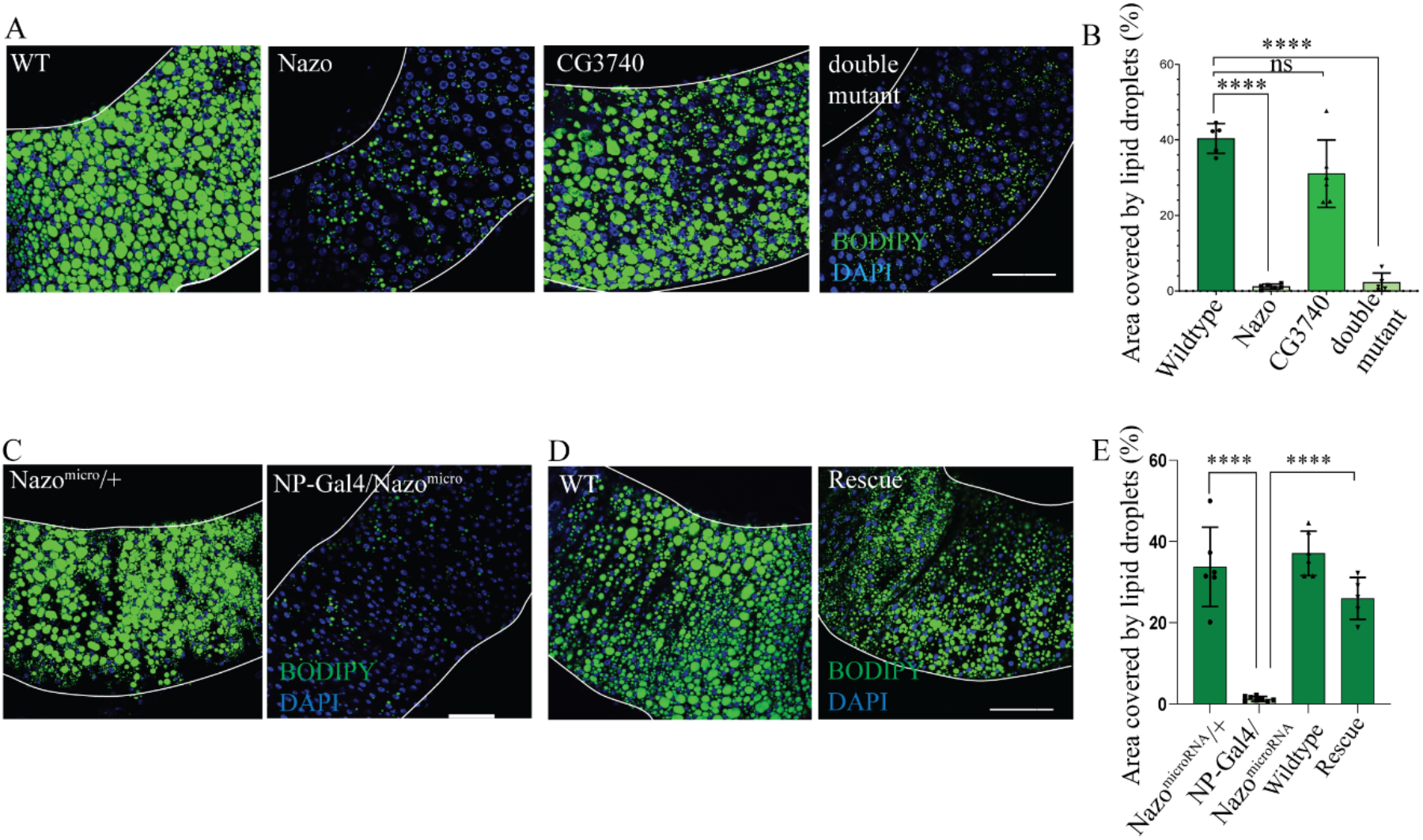
*nazo* mutants grown in axenic conditions also show gut lipid droplet depletion. Nazo is required in enterocytes for gut lipid homeostasis. (A) Guts from 20-day old flies raised in axenic conditions show reduced levels of lipid droplets in *nazo* and *nazo/CG3740* double mutants compared to wildtype flies (BODIPY™ 493/503 dye – green and DAPI – blue; Scale bar = 10µm) (B) Quantification of percentage area occupied by lipid droplets in the midgut R4 region of axenic flies of indicated genotype (N=20; Student’s t-test **** p-value =< 0.0001; ns =not significant)(C) Guts from 20-day old flies expressing Nazo microRNA under enterocyte-specific NP-Gal4 driver show reduced lipid droplets compared to control flies. (BODIPY™ 493/503 dye – green and DAPI – blue; Scale bar = 10µm) (D) Enterocyte-specific NP-Gal4 driven expression of UAS-Nazo transgene in nazo mutant background rescues the gut lipid droplet depletion phenotype. (BODIPY™ 493/503 dye – green and DAPI – blue; Scale bar = 10µm) (E) Quantification of the area covered by lipid droplets in the midgut R4 regions of indicated genotypes. (N=20; Student’s t-test **** p-value =< 0.0001)

### Drosophila Nazo is required in enterocytes for gut triglyceride homeostasis

The lipid depletion phenotype in *nazo* mutants can be due to a defect in feeding or absorption of fat in the gut. To assess the feeding behavior, we performed a well-established feeding assay ^32^, in which the flies are starved for two hours and subsequently provided food containing blue dye.

The total amount of food consumed in 24 hours was calculated using colorimetry for the blue dye. This assay did not reveal any defect in feeding behavior in the mutants (Supplementary Fig. 1B). To test whether the absorption of fatty acids into the gut was affected in the mutant alleles, we fed the adult flies with food containing fluorescently tagged oleic acid (BODIPY C16). 48 hours after feeding, the guts were fixed, and examined by fluorescence microscopy. There was no obvious difference in the levels of gut-incorporated fluorescent oleic acid between the wild-type control and the mutants suggesting that absorption of fatty acids was not defective in *nazo* mutants (Supplementary Fig. 1A). In addition, whole body glucose quantification did not reveal significant differences in overall glucose levels in WT vs mutant flies (Supplementary Fig. 1C).

To define the cell type in which nazo is required for the maintenance of gut lipid levels, we examined the effect of selective depletion of nazo using various cell type-specific Gal4 drivers. To this end, we generated transgenic lines harboring two microRNAs in tandem, targeting two distinct sequences in the *nazo* transcript ^33^. Depletion of nazo in enterocytes using the NP-Gal4 driver (Fig. 3C, E) resulted in the loss of lipid droplets in the gut at levels similar to the *nazo* null mutants, whereas neuron-specific knock down using the elav-Gal4 had no significant effects. Using an alternative approach, we assessed if NP-Gal4 driven enterocyte specific expression of Nazo transgene can rescue the nazo mutant phenotype. Indeed, expression of nazo in enterocytes leads to restoration of lipid droplets in the guts of *nazo* mutants (Fig. 3D, E). Collectively, these findings indicate that nazo is required in enterocytes for maintenance of gut lipid droplets, which is also consistent with the marked enrichment of nazo in the gut.

### *Drosophila c19orf12* mutants have diminished triglyceride stores and are sensitive to starvation

Most energy reserves in flies are in the form of lipids droplets in the fat body. Given the gut lipid depletion phenotype of *nazo* mutants, we measured the lipid droplet content of fat bodies in 40-day old adult males by BODIPY™ 493/503 staining (Fig. 4A, B). This revealed a significant reduction in fat body lipid droplet content in *nazo* mutants as well as *nazo*/CG3740 double mutants compared to WT flies. The phenotype is also present but less pronounced in younger flies. Ubiquitous expression of actin-Gal4 driven nazo microRNA also recapitulates this phenotype (Supplementary Fig. 2A, B). We also quantified the triglyceride levels from the whole bodies of the animals (Fig. 4C).

**Figure 4.**
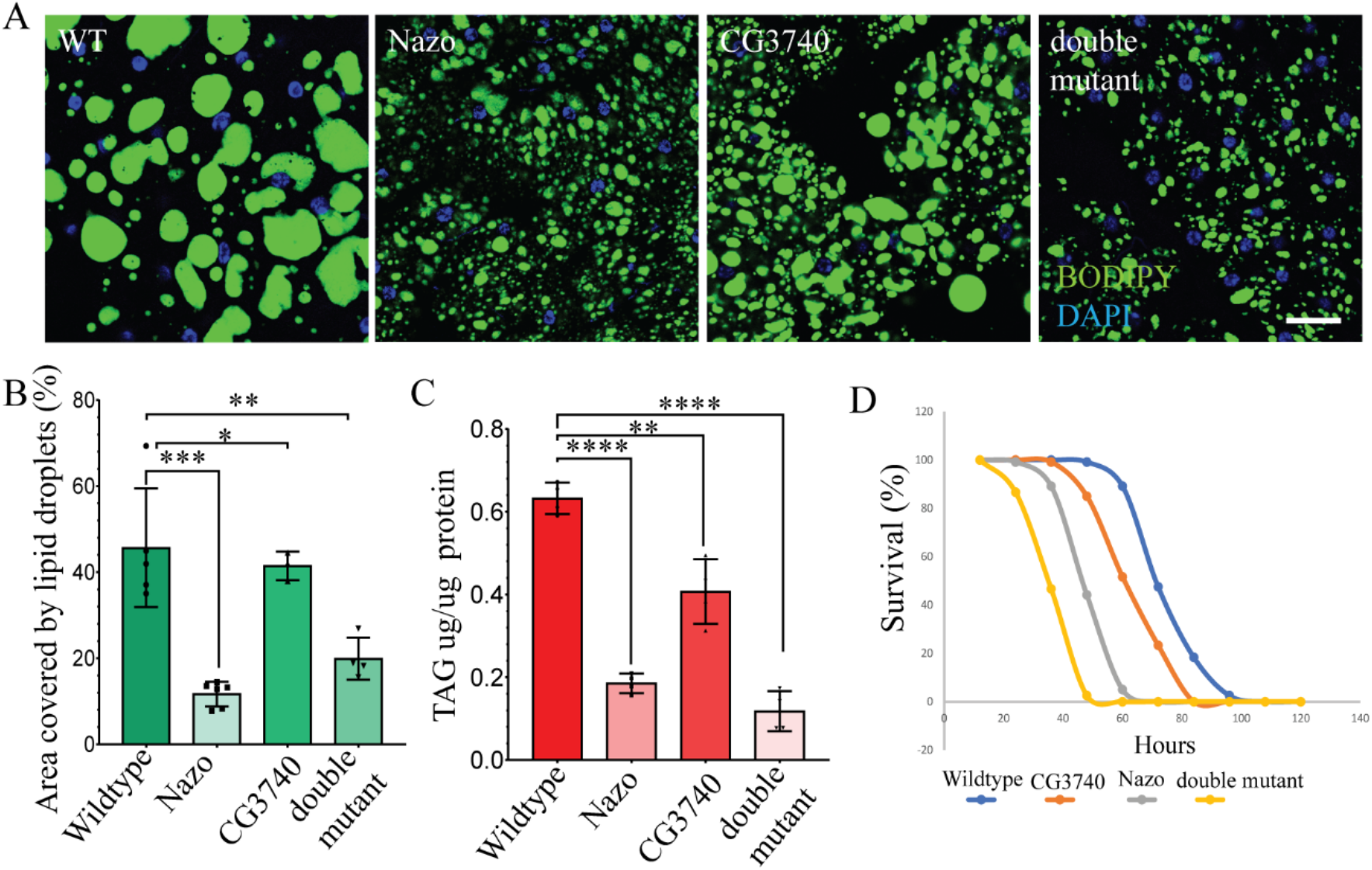
Flies lacking c19orf12 homologs are sensitive to starvation due to diminished fat body lipid droplets and whole-body triglyceride stores. (A) Confocal images of fat-body of 40-day old adults of indicated genotype, stained with BODIPY™ 493/503 dye (BODIPY™ 493/503 dye – green and DAPI – blue; Scale bar = 10µm) (B) Quantification of percentage area covered by lipid droplets in each fat body (N=20; Student’s t-test, p-value \*\*\**p* < 0.005) (C) Quantification of the TAG levels in whole body of 20-day old males. 8 males per genotype per biological replicates were used in quadruplets. (Student’s t-test **** p-value =< 0.0001) (D) Percentage of surviving flies of indicated genotypes under starvation conditions, counted every 12 hours (N=100)

In line with the previous findings, we observe significant reduction in the total triglyceride content in the *nazo* mutant and *nazo/CG3740* double mutants compared to wildtype flies (Fig. 4C). In these assays, *CG3740* mutants also show mild reduction in whole-body triglyceride content. Since *nazo* mutants show reduced lifespan along with diminished triglyceride levels, we asked whether these mutants are sensitive to starvation. To test this hypothesis, we subjected adult male flies to starvation stress (Fig. 4D) by transferring them to vials containing wet filter paper without any other nutrients. The percentage of dead flies were counted every 12 hours. This revealed that *nazo* mutants or *nazo/CG3740* double mutants are more sensitive towards starvation (Fig. 4D). In summary, these observations suggest that in addition to gut lipid depletion *nazo* mutants exhibit diminished fat body and whole-body triglycerides stores, which manifests as vulnerability to starvation.

### *nazo* mutants are sensitive to oxidative stress

The inability to sequester fatty acids in the lipid droplets results in oxidative stress ^34^. Considering the diminished triglyceride stores in *nazo* mutants, we decided to examine if these mutants are more sensitive to oxidative stress. To this end, WT and mutant flies were transferred to vials containing wet filter paper with 5% H_2_O_2_ and the percentage of dead flies was analyzed every 12 hours. Both *nazo* and *nazo/CG3740* double mutants revealed similarly increased sensitivity to H_2_O_2_ compared to wild type flies (Fig. 5A). Prompted by this finding, we examined the total levels of H_2_O_2_ in whole bodies of 20-day-old WT and *nazo* mutant males. We saw a considerable increase in overall H_2_O_2_ levels in *nazo* mutants, compared to wildtype control (Fig. 5B). There was another intriguing finding in these mutants. When raised on high fat diet (30% coconut oil) *nazo* as well as *nazo/CG3740* double mutants develop a dramatic 5-fold enlargement of their crops on average (Fig. 5C). Dye based gut motility assay revealed that this was due to diminished gut motility (data not shown). These findings raise the possibility that due to defects in triglyceride metabolism *nazo* mutants retain fat-rich food in their crops. Together, these data demonstrate various impacts of lipid dyshomeostasis in *nazo* mutants including increased sensitivity to oxidative stress.

**Figure 5.**
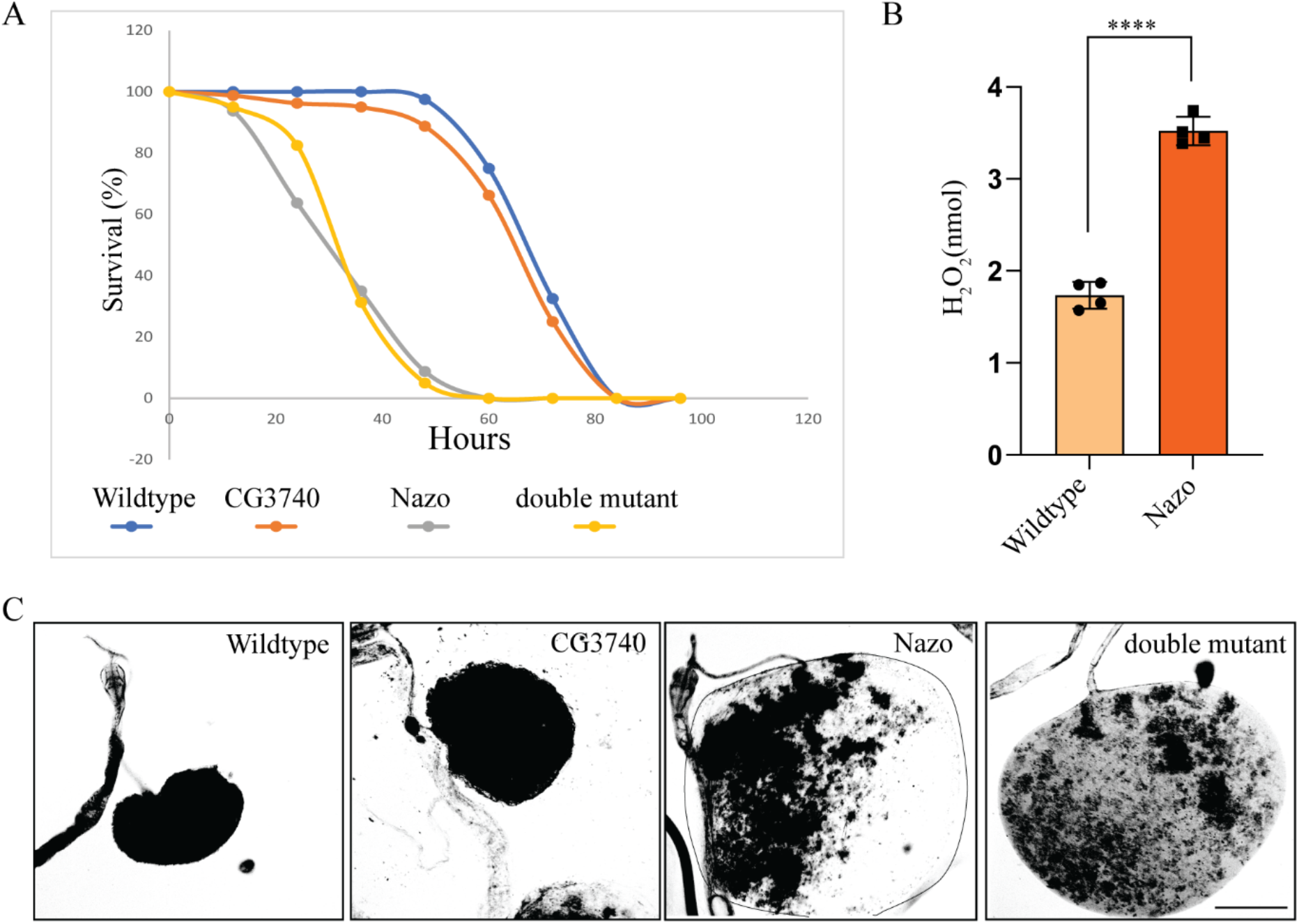
*nazo* mutants are sensitive to oxidative stress and show marked crop enlargement when raised on high fat diet. (A) Percentage of surviving flies of indicated genotypes raised on food supplemented with 5% H_2_O_2_, counted every 12 hours (N=80) (B) Whole body H_2_O_2_ quantification shows that *nazo* mutants have increased H_2_O_2_ levels relative to WT flies. 8 males per genotypes per biological replicates were used in quadruplets. (Student’s t-test **** p-value =< 0.0001) (C) Bright field images of the crops of 20-day old flies of indicated genotypes raised on high-fat diet for 3 days (N=20; Scale bar = 5µm)

### Nazo is an ER-lipid droplet contact site protein

To better understand the physiological role of *nazo* we examined its localization. As antibodies for Drosophila *nazo* are not available, we expressed HA-tagged nazo in S2 cells. Considering the triglyceride depletion phenotype of *nazo* mutants, we examined the localization of nazo with respect to lipid droplets. Therefore, the transfected S2 cells were treated with oleic acid to induce lipid droplet formation.

The cells were stained with antibodies against HA and Calnexin, an endoplasmic reticulum marker as well as the lipid dye BODIPY™ 493/503. Nazo localizes to discrete foci that overlap with endoplasmic reticulum. Interestingly, a significant proportion of the nazo-positive foci reside at the periphery of lipid droplets suggesting that they represent membrane contact sites between endoplasmic reticulum and lipid droplets (Fig. 6A). This finding is consistent with the role played by nazo in gut lipid homeostasis.

**Figure 6.**
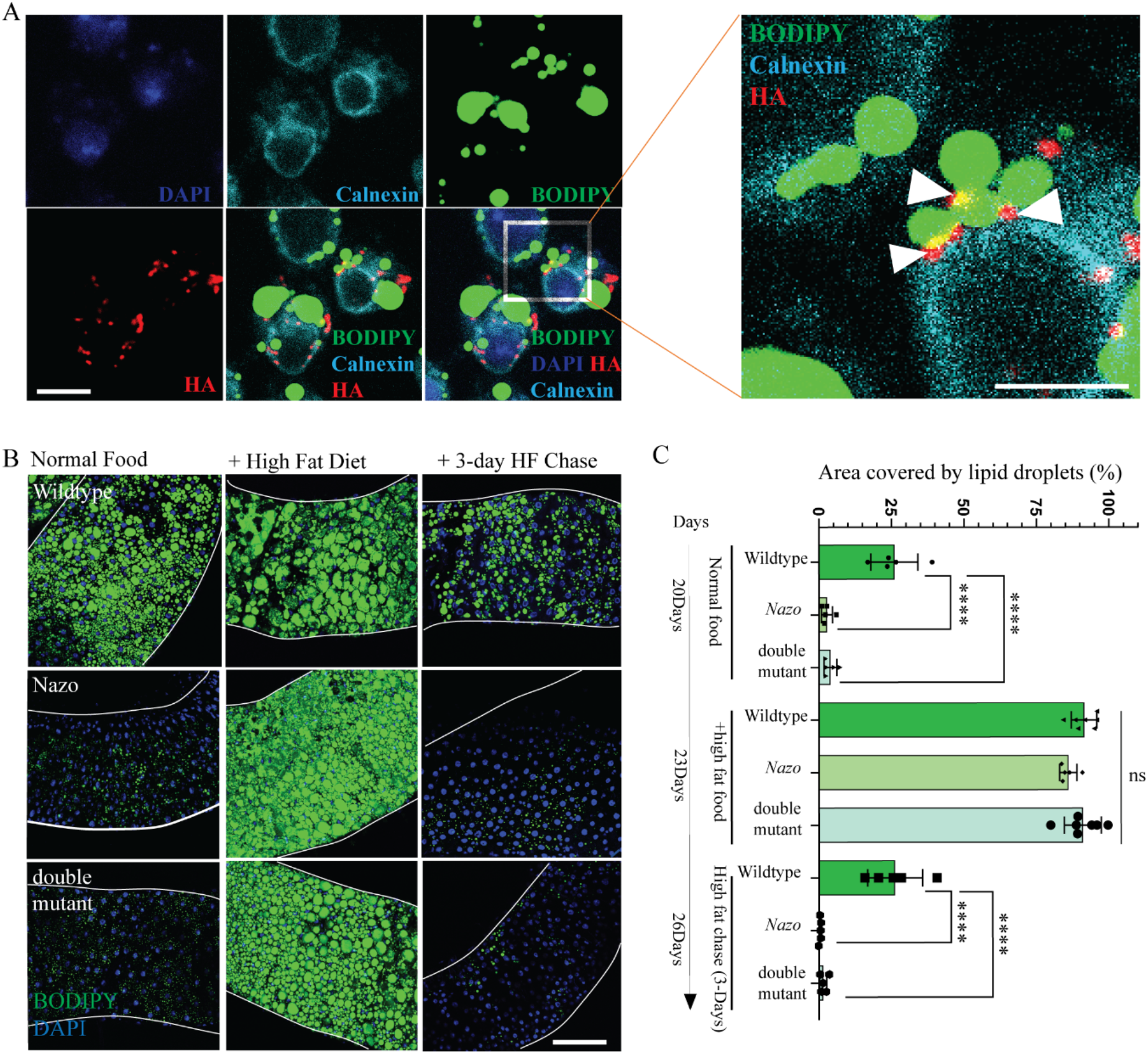
Nazo is an ER-lipid droplet contact site protein that is required for maintaining lipid droplet content on normal diet. (A) Confocal images of HA-Nazo expressing, oleic-acid treated S2 cells, stained with ER-Marker calnexin (cyan) and HA (red) antibodies, DAPI (blue) and BODIPY™ 493/503 dye (green), shows that Nazo localizes to ER-lipid droplet contact sites. (Scale bar =100µm) (B) Confocal images of BODIPY™ 493/503 stained guts from flies of indicated genotypes raised on normal diet or high-fat diet for 3 days or transferred to normal diet for 3 days after high-fat diet; high-fat diet rescues gut lipid droplet depletion in *nazo* and *nazo/CG3740* double mutants, but the phenotype reappears when flies are transferred back to normal diet (C) Quantification of the percentage area occupied by lipid droplets in indicated genotypes and conditions.

### The lipid droplet depletion in nazo mutants is due to dysregulated lipolysis

The lipid droplet depletion in *nazo* mutants can arise from decreased anabolism or increased catabolism of triglycerides. We first sought to address if fatty acids derived from triglycerides in the food can be properly absorbed and incorporated into the lipid droplets in nazo mutants. To test this possibility, we examined if high fat diet can rescue the *nazo* mutant phenotype. 20-day-old wild type and mutant flies were fed on a diet containing 30% coconut oil for 3 days and gut lipid droplet was examined by BODIPY™ 493/503 staining. While *nazo* mutants on normal diet demonstrated significant gut lipid droplet depletion, mutants placed on high fat diet showed restoration of gut lipid droplet content (Fig. 6B, C). We next asked if gut lipid droplets can be maintained in these flies if they are transferred back to normal food from high-fat diet? Interestingly, three days after transfer to normal diet *nazo* mutants redevelop lipid depletion phenotype (Fig. 6B, C). Overall, these findings suggest that fatty acids from diet can be properly incorporated into lipid droplets in *nazo* mutants but likely due to increased catabolism these mutants are not able to maintain their triglyceride content.

To examine the impact of lipolysis on *nazo* mutant phenotype, we asked if depletion of lipases can suppress the gut lipid droplet depletion in *nazo* mutants. We tested the roles of three different lipases including Brummer, Dob and CG5966 in this context by RNAi mediated depletion. We generated a line *CG3740/FM6; Actin-Gal4::Nazo* ^*microRNA*^*/Cyo* that harbors the CG3740 mutation and lacks nazo due to *actin-Gal4* driven ubiquitous expression of Nazo microRNA. This line shows the same level of gut lipid droplet depletion as the *nazo null* mutant. This line was crossed with *Brummer*^*RNAi*^, *Dob*^*RNAi*^, and *CG5966*^*RNAi*^ lines to examine the effect of concurrent depletion of these lipases on *nazo* mutant phenotype. Interestingly, loss of Brummer but not Dob or CG5966, leads to rescue of the *nazo* mutant phenotype (Fig. 7A, B and data not shown).

**Figure 7.**
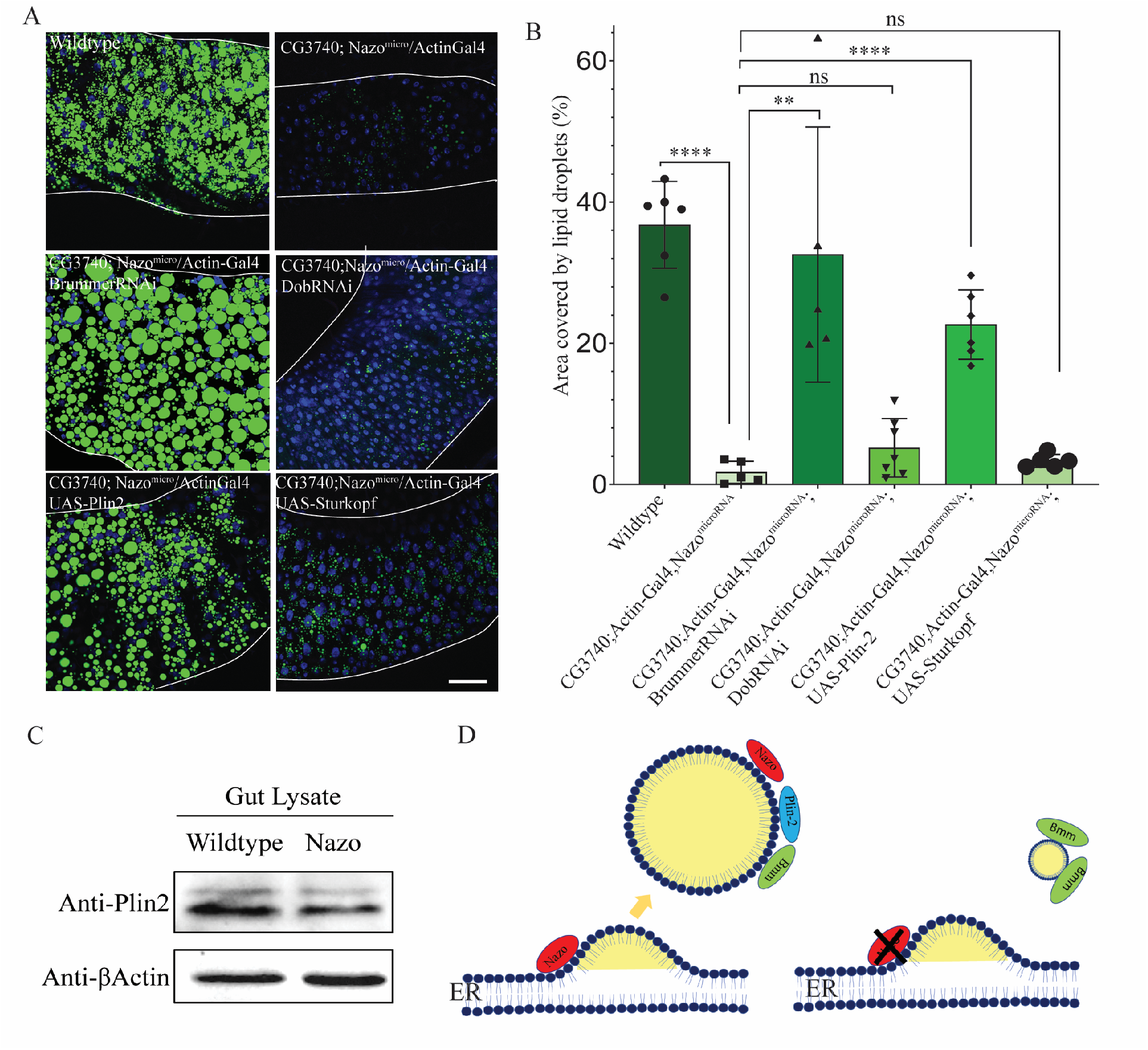
The lipid droplet phenotype in Nazo mutant is due to dysregulated lipolysis. (A) Confocal images of guts from 20-day old flies of indicated genotypes stained with BODIPY™ 493/503 dye (BODIPY™ 493/503 dye – green and DAPI – blue; Scale bar = 10µm); Concurrent Knockdown of the lipase Brummer but not Dob can rescue the reduced gut lipid droplet phenotype resulting from nazo depletion. Similarly, overexpression of Perilipin-2, a regulator of Brummer, but not Sturkopf (another lipid metabolism gene) rescues the nazo depletion phenotype (B) Quantification of the percentage area occupied by lipid droplets midguts of flies of indicated genotypes (C) Immunoblot showing reduced expression of perilipin -2 in gut lysate from *nazo* mutants relative to wild type flies (D) Model for the role of Nazo in lipid homeostasis. Nazo is a membrane contact site protein that maintains lipid droplets by restraining the lipase activity of Brummer.

One of the critical regulators of Brummer is Perilipin-2, a lipid droplet protein, which maintains lipid droplets by antagonizing Brummer ^35,36^. Like Nazo, depletion of Perilipin-2 leads to diminished fat stores. Therefore, we decided to check if the level of Perilipin-2 is altered in nazo mutants. Western blot analysis of gut lysates revealed that Perilipin-2 levels are lower in nazo mutants relative to WT flies (Fig. 7C). This prompted us to test if Perilipin-2 overexpression can rescue the lipid depletion phenotype of *nazo* mutants.

As discussed earlier, microRNA mediated depletion of nazo results in loss of lipid droplet in the gut. Co-expression of Perilipin-2 in this setting, however, leads to suppression of this phenotype. We also tested the impact of overexpression of Sturkopf, another lipid droplet associated protein in this context. Unlike Perilipin-2, however, Sturkopf overexpression could not suppress the *nazo* mutant phenotype (Fig. 7A, B). Overall, these experiments provide a plausible explanation for lipid droplet defect in *nazo* mutants. These mutants exhibit diminished Perilipin 2, which in turn may result in increased lipolysis due to overactivity of the lipase Brummer.

In summary, we show that the Drosophila c19orf12 homolog – Nazo is a membrane contact site protein, which maintains gut lipid droplets by regulating lipolysis. Given the role of lipid dyshomeostasis in neurodegeneration, our findings provide the foundation for future studies on the role of mammalian c19orf12 in lipid homeostasis and neurodegeneration.

## DISCUSSION

Multiple lines of evidence point to a role for c19orf12 in lipid homeostasis. First, human adipose tissue and other adipocyte-rich organs such as breast and liver show striking enrichment of c19orf12 (Supplemental fig. 3, spliced from RNA expression consensus dataset from human protein atlas). Second, ablation of the fly c19orf12 homologs results in differential expression of multiple lipid metabolism genes in guts. Finally, and most importantly, flies lacking the c19orf12 homolog – Nazo exhibit a robust and completely penetrant lipid droplet depletion phenotype in their guts. Nazo plays a cell autonomous role in enterocytes to maintain the gut lipid droplet pool. This phenotype is not exacerbated by the added absence of CG3740, suggesting that Nazo is the primary player in this context. It is intriguing that even though CG3740 is expressed more widely than Nazo, its impact on the overall fitness of the organism (as assessed by locomotor and viability assays) is significantly less pronounced than Nazo. The function of CG3740 is still unclear and future studies are needed to investigate its role in neuronal and nonneuronal tissues. In summary, the expression pattern of c19orf12 in humans and the physiological role of Nazo in *Drosophila* is suggestive of a conserved role for this protein in lipid homeostasis.

What are the functional consequences of gut triglyceride depletion observed in *nazo* mutants? Disruption of Nazo has profound impact on the health of animals as evidenced by reduced life span and locomotor deficits in these mutants. Triglyceride stores are particularly important for the sustenance of animals during phases of starvation. Thus, as expected, *nazo* mutants are highly vulnerable to starvation. In addition to fulfilling the energetic needs of the animal, lipid droplets also play a protective role against reactive oxygen species by sequestering fatty acids into triglycerides and preventing aberrant lipid peroxidation. In concordance with their lipid depletion phenotype, ablation of Nazo leads to increased sensitivity to hydrogen peroxide induced oxidative stress. Thus, compromised lipid metabolism in *nazo* mutants has significant implications for the sustenance of animals.

Nazo was identified as one of the genes upregulated in response to viral infection in a NF-κB dependent manner ^30^. Innate immune response is intricately linked to metabolism and multiple studies have shown that bacterial infection and innate immune response effect gut and fat body triglyceride levels ^19-21^. While it is plausible that Nazo serves as a link between immune response and metabolism by modulating triglyceride homeostasis, our findings suggest that Nazo governs lipid homeostasis under physiological conditions independent of its possible role in immune response. We observe lipid droplet depletion in *nazo* mutants in normal rearing conditions in the absence of extraneous bacterial infection. More importantly, this phenotype is observed in axenic flies, which are raised in the presence of antibiotics and lack gut microbiota indicating the gut commensal bacteria are not responsible for this phenotype. Interestingly, two studies have recently shown that NF-κB and sting influence triglyceride metabolism in an immune response-independent fashion. While *NF-κB (relish)* mutants exhibit a rapid decline in fat body triglyceride stores on starvation, *sting* mutants show diminished triglyceride stores in fat body even on normal diet ^22,23^. These studies prompted us to examine the gut lipid droplet levels in NF-κB mutants or sting RNAi flies. Unlike *nazo* mutants, NF-κB mutants or sting RNAi flies did not reveal diminished gut lipid droplets under normal rearing conditions. The latter observation is consistent with the report from Akhmetova et al who did not observe reduced gut lipid droplet content in *sting* mutants ^22^. Overall, our findings suggest that the lipid droplet depletion observed in nazo mutants is likely not due to disruption of immune response or dysregulation of NF-κB pathway.

The subcellular localization of nazo is further suggestive of the fact that it plays a direct role in lipid metabolism. In S2 cells treated with oleic acid nazo localizes to foci in endoplasmic reticulum that lie in close proximity to lipid droplets suggesting that they belong to membrane contact sites between lipid droplets and endoplasmic reticulum. The localization of human c19orf12 has not been examined under conditions of lipid droplet induction. However, in cultured cells grown on normal media, human c19orf12 localizes to endoplasmic reticulum and mitochondria, raising the possibility that under basal conditions it may be a component of the ER-mitochondria contact site ^27^. Proteins like Seipin and Vps13a are known to localize to contact sites of various types including ER-mitochondria and ER-lipid droplet contact sites ^37,38^. Thus, it needs to be explored if human or fly c19orf12 homologs localize to and function at multiple distinct membrane contact sites including ER-LD contact sites.

Many of our observations support that increased lipolysis is responsible for lipid droplet depletion in *nazo* mutants. First, *nazo* mutant phenotype can be suppressed by feeding high fat diet to flies. When these flies are transferred back to normal diet, their lipid stores are depleted within a few days. Second, simultaneous depletion of the lipase – Brummer, rescues the gut lipid depletion observed in response to suppression of Nazo. Third, Perilipin 2, an inhibitor of Brummer is diminished in nazo mutants suggesting that Brummer overactivity may be responsible for triglyceride depletion in nazo mutants. Finally, overexpression of Perilipin 2 rescues the nazo mutant phenotype. Overall, these findings suggest that nazo is an ER-lipid droplet contact site protein that regulates lipolysis and maintains gut triglyceride levels through its effect on Brummer-Perilipin 2 axis. Given the localization of Nazo to ER-lipid droplet contact sites, it is interesting that Brummer localizes to discrete foci on lipid droplets ^13^. It needs to be investigated if these foci represent ER-lipid droplet contact sites and whether Brummer’s localization is affected in *nazo* mutants. The role of ER-lipid droplet contact sites in the biogenesis of lipid droplets is well established. Studies in yeast and cell culture systems have demonstrated that in the absence of ER-LD contact site proteins like Seipin, lipid droplets show significant morphological abnormalities ^11,12^. What is less well understood is the role of ER-LD contact sites in the maintenance of lipid droplets by limiting lipolysis and lipophagy. In mammals, Seipin deficiency leads to increased lipolysis and marked loss of adipocytes, which can be partially rescued by the suppression of the lipase ATGL that is homologous to *Drosophila* Brummer ^39,40^. Thus, abrogation of ER-LD contact sites not only disrupts LD biogenesis but also promotes triglyceride catabolism. The morphologically abnormal lipid droplets generated in the absence of Seipin also show a remarkable alteration in their proteome ^11^. Thus, it is possible that unrestrained lipolysis in *nazo* mutants stems from defects in ER-LD contact sites leading to diminished incorporation of lipase inhibitor proteins like Perilipin-2, during LD biogenesis.

Mutations in human c19orf12 gene cause NBIA characterized by neuronal loss that is most prominent in basal ganglia ^25^. Though extensive iron accumulation is present in the basal ganglia of these patients it is unclear whether iron accumulation is the primary cause of neurodegeneration or if it is an epiphenomenon. In NBIA patients, neuronal dysfunction/loss and functional abnormalities are observed in parts of brain other than basal ganglia where iron accumulation is not prominent. Furthermore, only two of the NBIA-mutated genes are directly involved in iron metabolism whereas the others function in diverse processes including lipid homeostasis, mitochondrial function and autophagolysosomal degradation ^26^. Though, it has been challenging to come up with a unifying mechanism for this disease, there is substantial evidence to suggest that disruption of lipid metabolism plays a key role in the pathogenesis of this disease. Two of the NBIA-mutated genes PLA2G6 and FAAH are directly involved in lipid metabolism. Two other genes – COASY and Pantothenate kinase are crucial components of the Coenzyme A biosynthesis pathway, a critical regulator of various aspects of lipid metabolism ^26^. In continuation with this theme, our study provides the first evidence for the involvement of the fly c19orf12 homolog – Nazo in lipid homeostasis. Our findings provide a foundation for future studies, seeking to decipher the role of mammalian c19orf12 in neuronal and nonneuronal tissues, especially in the context of lipid biology.

An interesting feature of two of the NBIA subtypes caused by *c19orf12* and *PLA2G6* mutations is the dramatic accumulation of synuclein-positive inclusions in various parts of brain. This suggests that there may be mechanistic similarities between NBIA and sporadic Parkinson’s disease and studies on these proteins may inform us regarding the mechanism of other synucleinopathies. It is being increasingly appreciated that lipid dyshomeostasis plays an important role in synuclein aggregation. To list just a few observations, inhibition of Stearoyl-CoA desaturase, an enzyme required for synthesis of monosaturated fatty acids can inhibit synuclein accumulation in vitro and in vivo ^41-43^. Furthermore, Mori et al have demonstrated that mutations in the phospholipase PLA2G6 leads to shortening of phospholipid acyl chains which in turn promotes accumulation of α-synuclein ^44^. Thus, it is tempting to speculate that the synuclein accumulation observed in human patients with c19orf12 mutants is due to defects in lipid metabolism.

In summary, this study establishes a role for the c19orf12 homolog – Nazo in lipid homeostasis and provides novel insights into the mechanism of NBIA. Our findings will pave the way for exploring the connection between lipid dyshomeostasis, neurodegeneration and synuclein accumulation.

## METHODS

### Fly husbandry and genetics

The flies were reared in standard cornmeal/agar medium supplemented with yeast at 25°C with 60 ± 5% relative humidity and 12h light /dark cycles unless specified. The following strains were obtained from Bloomington Drosophila Stock Center: *w*^*1118*^, *Actin-Gal4* (25374), *NP-Gal4, elav-Gal4, Brummer*^*RNAi*^ (25926), *Dob*^*RNAi*^ (65925), *CG5966* ^*RNAi*^ and *UAS-Sturkopf* (91394). *UAS-Perilipin2* ^13^ was a kind gift from Dr. Ronald Kuhnlein.

The following lines were generated for this study *CG3740 null, nazo (CG11671) null, UAS-CG11671*^*myc*^ and *pUAST-CG11671*^*microRNA*^.

CG3740 and CG11671(nazo) knockout allele was created by CRISPR/Cas9-mediated gene editing according to a published procedure ^45^. Briefly, we replaced part of the coding sequence of these genes with DsRed through homology-mediated repair. Two different guide RNAs were used for each of these genes. Sequences flanking the guide RNA target sites were amplified from genomic DNA and introduced into donor plasmids to facilitate homology-directed repair.

The following primer sequences were used for CG3740 guide RNAs:

5’-Guide RNA

> Sense oligo: 5’- CTTCGGAGGCGATAGCCATCGTAG -3’
>
> Antisense oligo: 5’- AAACCTACGATGGCTATCGCCTCC -3’

3’-Guide RNA

> Sense oligo: 5’- CTTCGCGGTTGGCGGGGCCGCCGG -3’
>
> Antisense oligo: 5’- AAACCCGGCGGCCCCGCCAACCGC -3’

The following primers were used to generate homology arms for CG3740:

5’-Homology arm

> Forward: 5’- ATATCACCTGCATATTCGCCAGGTACACCCATACACTAG -3’
>
> Reverse: 5’- ATATCACCTGCATATCTACCGATGGCTATCGCCTCCATC -3’

3’-Homology arm

> Forward: 5’- ATATGCTCTTCATATCGGCGGAATCGCCGCCTACAAGA -3’
>
> Reverse: 5’- ATATGCTCTTCAGACTAGGTTATTCCATTACCATTCGG -3’

The following primer sequences were used for CG11671 guide RNAs:

5’-Guide RNA

> Sense oligo: 5’- CTTCGTAACTCCCCTCCACAGTGG -3’
>
> Antisense oligo: 5’- AAACCCACTGTGGAGGGGAGTTAC -3’

3’-Guide RNA

> Sense oligo: 5’- CTTCGGGAATGACCATCGTTGATT -3’
>
> Antisense oligo: 5’- AAACAATCAACGATGGTCATTCCC-3’

The following primers were used to generate homology arms for CG11671:

5’-Homology arm

> Forward: 5’- ATATCACCTGCATATTCGCACTCGTGCCAACACTTCTTG -3’
>
> Reverse: 5’- ATATCACCTGCATATCTACCTGTGGAGGGGAGTTACTCG -3’

3’-Homology arm

> Forward: 5’- ATATGCTCTTCATAT<u>ATT AG</u>GCATTAATAGAAATA -3’
>
> Reverse: 5’- ATATGCTCTTCAGACTCACTATGTTCTTATAAAAC -3’

Both 5’ and 3’ homology arms were then cloned into the pHD-DsRed-attP vector containing the eye-specific 3xP3 promoter fused with DsRed and the resulting construct was microinjected into Cas9-expressing embryos by a commercial service (Rainbow Transgenic Flies Inc.). Flies bearing the *CG3740 or CG11671* deletion were identified by screening the offspring of injected adults for expression of red fluorescence in the eye. The deletion mutants were confirmed by PCR and sequencing, using primers against a dsRed sequence and adjacent CG3740/CG11671 genomics sequences. Subsequently, DsRed markers inserted at the deletion sites were removed by crossing the mutants with hsCre flies and maintaining the progenies at 25°C.

*CG11671* microRNA constructs targeting the sequences TTCAGCCATCTCAGAGATAATC and GGTCATTTTGAACGATT TGACG in the nazo transcripts were generated as described ^33^. Briefly, while keeping the stem-loop backbone and surrounding sequences unaltered, the *Drosophila* Mir6.1 gene-targeting microRNA sequence was replaced with the 22 bps complementary to the *nazo* transcript. Two microRNA constructs targeting distinct *nazo* sequences were inserted in tandem into the pUAST vector and the resulting plasmid was used to generate *UAS-Nazo*^*microRNA*^ transgenic flies. For rescue experiments, the *Drosophila* Nazo transgene was cloned into pUAST vector along with a carboxy-terminal 3 X MYC tag.

Unless specified, all experiments were performed on 20-day-old males from different genotypes.

### Life span assay

For life span analysis 60 males for each genotype were transferred to fresh food (20 flies per vial) every 2-3 days and dead flies were counted.

### Climbing assay

Climbing behavior was examined by Rapid Iterative Negative Geotaxis (RING) assay at 20 days age, according to a previously published protocol ^46^. Briefly, flies were transferred into plastic vials and six vials at a time were loaded onto the RING apparatus (10 flies per vial). The vials were tapped down to initiate the climbing response and the height climbed by each fly after 4 s was recorded. The climbing assay was repeated three times and the average height climbed in the three trials was calculated.

### RNA seq analysis

Total RNA was isolated from triplicates of 20 freshly dissected guts of 20-day old WT and *CG3740/nazo* mutant males using the RNeasy Plus Mini Kit (Qiagen, Valencia, CA) per manufacturer’s recommendations. RNA-seq analysis was performed at UR Genomics Research Center. The total RNA concentration was determined with the NanopDrop 1000 spectrophotometer (NanoDrop, Wilmington, DE) and RNA quality assessed with the Agilent Bioanalyzer (Agilent, Santa Clara, CA). The TruSeq Stranded mRNA Sample Preparation Kit (Illumina, San Diego, CA) was used for next generation sequencing library construction per manufacturer’s protocols. Briefly, mRNA was purified from 200ng total RNA with oligo-dT magnetic beads and fragmented. First-strand cDNA synthesis was performed with random hexamer priming followed by second-strand cDNA synthesis using dUTP incorporation for strand marking. End repair and 3` adenylation was then performed on the double stranded cDNA. Illumina adaptors were ligated to both ends of the cDNA, purified by gel electrophoresis and amplified with PCR primers specific to the adaptor sequences to generate amplified libraries of approximately 200-500bp in size. The amplified libraries were hybridized to the Illumina flow cell and single end reads of 75nt were generated for each sample using Illumina’s NextSeq550 sequencer.

Analysis methods: Raw reads generated from the Illumina basecalls were demultiplexed using bcl2fastq version 2.19.0. Quality filtering and adapter removal are performed using FastP version 0.20.0 with the following paramters: ““--length_required 35 --cut_front_window_size 1 --cut_front_mean_quality 13 -- cut_front --cut_tail_window_size 1 --cut_tail_mean_quality 13 --cut_tail -y -r”. Processed/cleaned reads were then mapped to the D. melanogaster reference genome (BDGP6.28) using STAR_2.7.0f with the following parameters: “—twopass Mode Basic --runMode alignReads --outSAMtype BAM SortedByCoordinate – outSAMstrandField intronMotif --outFilterIntronMotifs RemoveNoncanonical – outReads UnmappedFastx”. Genelevel read quantification was derived using the subread-1.6.4 package (featureCounts) with a GTF annotation file and the following parameters: “-s 2 -t exon -g gene_name”. Differential expression analysis was performed using DESeq2-1.22.1 with a P-value threshold of 0.05 within R version 3.5.1 (https://www.R-project.org/). A PCA plot was created within R using the pcaExplorer to measure sample expression variance. Heatmaps were generated using the pheatmap package were given the rLog transformed expression values. Gene ontology analyses were performed using the EnrichR package.

### Starvation and Oxidative stress sensitivity assay

For starvation analysis, 100 flies were transferred from normal food to vials containing Whatman filter paper soaked with PBS (20 flies per vial). Fresh PBS was added every 24 hours to prevent drying. Dead flies were counted every 12 hours.

For Oxidative stress 80 flies were transferred from normal food to vials containing 5% H_2_0_2_ (20 flies per vial). Dead flies were counted every 12 hours.

### Generation of axenic flies

Flies were reared in axenic conditions as described previously ^47^. Briefly, embryos were washed in 2.5% chlorine (50% bleach) for 2 minutes followed by 2-minute wash in 70% alcohol and 2-minute wash in distilled water. The sterile embryos were transferred to food containing antibiotics (Tetracycline, chloramphenicol, ampicillin, and erythromycin). Axenic flies were transferred to fresh food supplemented with antibiotics every 2 days, in triplicates. Axenic conditions were tested by grinding the flies in PBS and spreading the suspension on LB, MRS or Mannitol plates at 37°C or 30°C. Absence of bacterial colonies indicated that the flies were devoid of gut microbes.

### BODIPY™ 493/503 Staining

Appropriately aged guts were dissected in PBS followed by fixation in 4% paraformaldehyde in PBS for 30 mins. The guts were incubated in BODIPY™ 493/503 (1ug/ml, Thermo fisher Scientific) for 30 minutes in dark with DAPI. The guts were washed with PBS for 30 mins and then mounted on slides in 70% glycerol. The guts were imaged immediately by confocal microscopy. For BODIPY™ 493/503 staining of fat bodies, adult carcasses were filleted in PBS, fixed in 4% paraformaldehyde and incubated in BODIPY™ 493/503 for 30 minutes.

### Triglyceride Quantification

TAG quantification was performed using Triglyceride Quantification kit (Sigma-Aldrich). For TAG quantification of whole body, 8 males were used in triplicates for each sample. Triglyceride standard solution and glycerol standard solution were used as standards. Total protein level in the samples were determined by Pierce Bradford Assay kit (Thermo Scientific).

### Glucose quantification

Glucose quantification was performed using glucose assay kit (Sigma-Aldrich) from whole bodies of triplicates of 8 males. Total protein level in the samples were determined by Pierce Bradford Assay kit (Thermo Scientific) and used for normalization.

### Hydrogen Peroxide quantification

H_2_O_2_ quantification was performed on triplicates of 8 adult males using Amplex Red Hydrogen Peroxide assay kit (Invitrogen) according to manufacturer’s instructions. Serial dilution of H_2_O_2_ was used to generate a standard curve.

### Cell Culture and Transfection

Drosophila S2 cells were purchased form Drosophila Genomics Resource center and cultured in Shields and Sand M3 media (Sigma-Aldrich) supplemented with 10% Insect Media Supplement (Sigma-Aldrich). pAc5.1A-HA-CG11671 was transfected into the S2 cells using Lipofectamine 3000 (Invitrogen). Cells were grown in an incubator at 25°C.

### Immunostaining

S2 cells transfected with HA-tagged CG11671 were fixed in 4% paraformaldehyde and washed 3 times in PBS. After incubation in blocking solution (PBS + 0.2% Triton X-100 + 3% BSA) for 1 hr, cells were stained overnight with antibodies against HA (rabbit polyclonal, 1:1000, Invitrogen) and Calnexin (mouse monoclonal, 1:20, DSHB) in blocking solution. Subsequently, they were stained with secondary antibodies (1:500, anti-rabbit Alexa Flour™ 546 and anti-mouse Alexa Flour™ 647) for 2 hours. The guts were next stained with BODIPY™ 493/503 and DAPI for 30 mins. Imaging was performed by confocal microscopy.

### High fat diet experiments

20-day old males were transferred to vials containing food supplemented with 30% coconut oil for 3 days. For High fat diet chase experiments, after 3 days on high fat diet, flies were transferred back to normal food. After 3 days in normal food, gut lipid droplet content was evaluated by BODIPY™ 493/503 staining.

### Western blot

Briefly, 10 guts per samples were lysed in 2 X SDS-loading buffer and protein was quantified using Pierce Detergent compatible Bradford Assay kit (Thermo Scientific). The lysates were separated by SDS-PAGE and transferred to Nitrocellulose membrane. The membrane was blocked with 5% non-fat dry milk in TBST and incubated with primary antibodies against Perilipin-2 (1:5000 gift from Dr. Ronal Kuhnlein) ^13^ and Actin (1:1000, Cell Signaling). The western blot images were captured by a BIO-RAD workstation.

### Consumption-excretion food assay

As previously described ^32^, feeder caps containing dyed (Blue-1) food media (2%) were placed on vials containing 15 adult males, which were starved for 2 hours. The flies were allowed to consume food for 24 hours. 24 hours post feeding, the flies were homogenized in water to collect the dye inside (Internal) the flies. The dye excreted by the animals were collected from the walls of the vials using water (External). The absorbance of the internal and external dye was measured by a spectrophotometer. Absorbance values were converted to volume of medium consumed by interpolation of the standard curve for pure dye. Flies which were fed with food without the dye were used for background absorbance.

### Imaging and signal analysis

Confocal images were collected using a Leica SP5 confocal system and processed using the Leica-LAS-X software, analyzed using the Fiji/ImageJ software package and were assembled using Adobe illustrator. All the gut images were captured from the R4 region of the midgut. For the quantification of the BODIPY signal, images were analyzed with the Fiji/ImageJ software package. Average pixel intensity in each gut/ fat body was calculated and normalized by subtracting the average pixel intensity in the background region. All the box plots were analyzed using GraphPad Prism 9.0 software.

### Lipid incorporation assay

For the incorporation of fluorescently labelled lipid in the lipid droplets flies were fed with yeast paste containing 1% sucrose and BODIPY F16 (Invitrogen) for 2 days. The guts were dissected in 1XPBS and fixed in 4% paraformaldehyde for 30 minutes. After fixation the guts were mounted in 70% glycerol and imaged by confocal microscopy.

### Statistics

All p-values were calculated using the Student’s t-test with unpaired samples. All error bars represent standard error of mean. Exact values of all *n*’s can be found in Figure legends.

### Materials availability

Transgenic flies and plasmids generated in this study will be available upon request.

## AUTHOR CONTRIBUTIONS

R.B., S.P., S.L., K.P., M.G. and N.M. performed experiments and analyzed data. R.B. provided project conceptualization, study design and supervised the project. R.B. and S.P. wrote the manuscript with input from all authors. R.B. and L.J.P. acquired research funding.

## ACKNOWLEDGEMENTS

This work was supported by National Institute of Health (T32 grant to RB as a fellow) and (R01 grant to LJP).

---

**Supplementary Figure 1.**
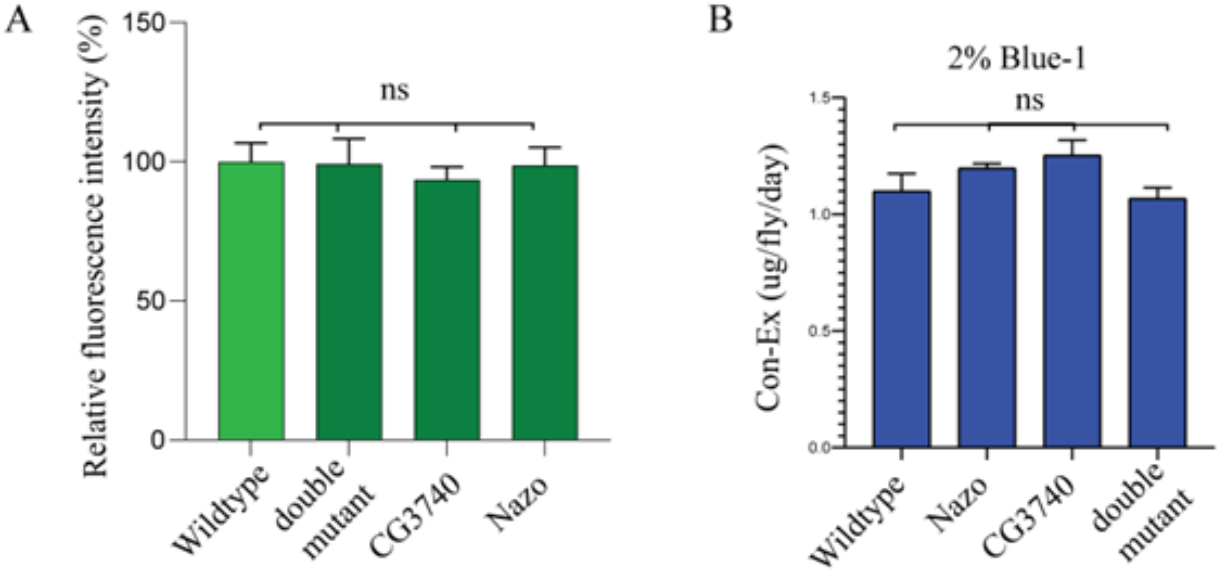
Nazo mutants do not show defects in feeding or absorption of fluorescent labeled oleic acid. (A) Quantification of gut fluorescence in flies of indicated genotypes fed with fluorescent oleic acid. (N=20, Student t-test, p-value ns= not significant) (B) Quantification of the amount of food consumed by adult males in 24-hour period laced with Blue-1 food dye color. (N=20, Student t-test p-value ns = not significant) (C) Quantification of whole-body glucose from males of indicated genotypes. 8 males per genotype per biological replicates were used in triplets. (Student t-test p-value ns = not significant)

**Supplementary Figure 2.**
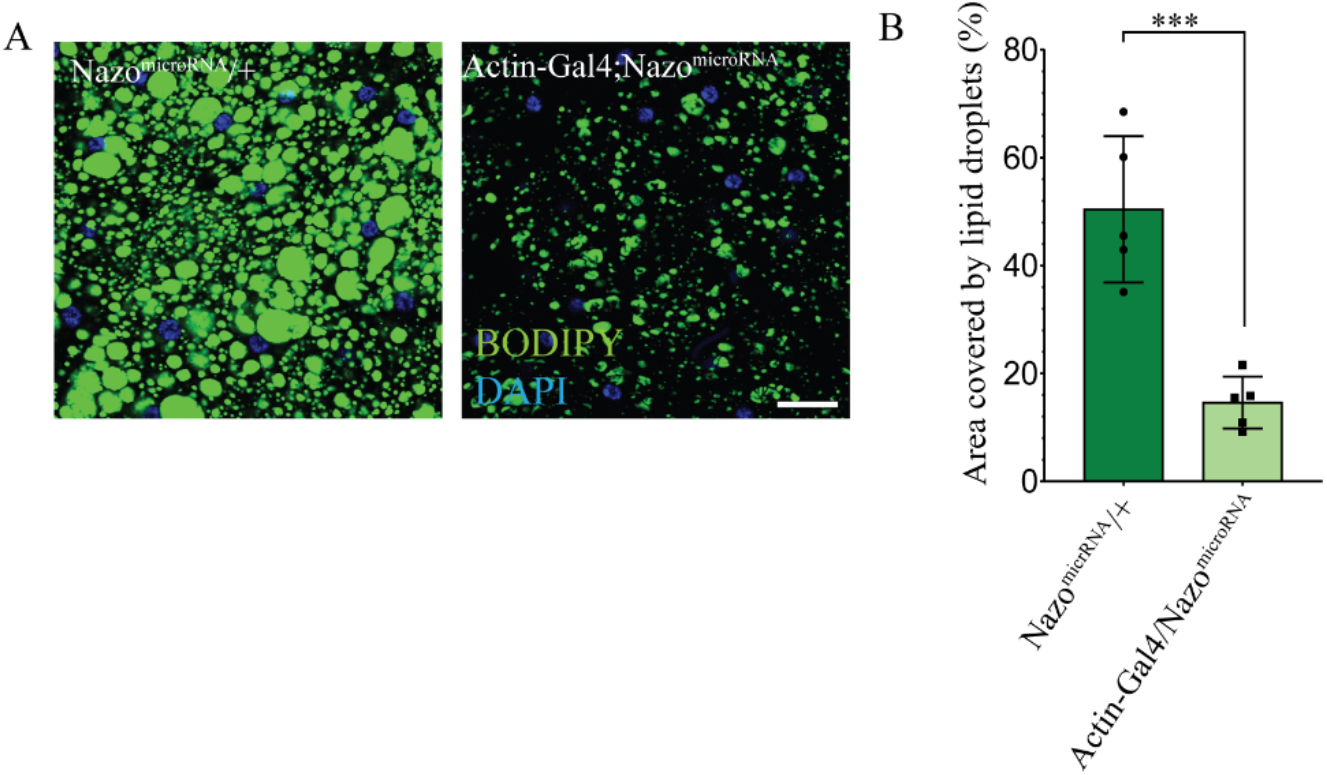
Micro-RNA mediated depletion of Nazo leads to diminished lipid droplets in fat bodies of adult flies. (A) Confocal image of fat body stained with BODIPY™ 493/503 dye from adult males shows that expression of Nazo^microRNA^ under the ubiquitous driver Actin-Gal4 leads to diminished lipid droplets in fat body relative to control flies (BODIPY™ 493/503 dye – green and DAPI – blue; Scale bar = 10µm))(B) Quantification of the percentage area occupied by lipid droplets in the fat bodies of indicated genotypes (N=20; Student’s t-test *** p-value =< 0.005)

**Supplemental figure 3.**
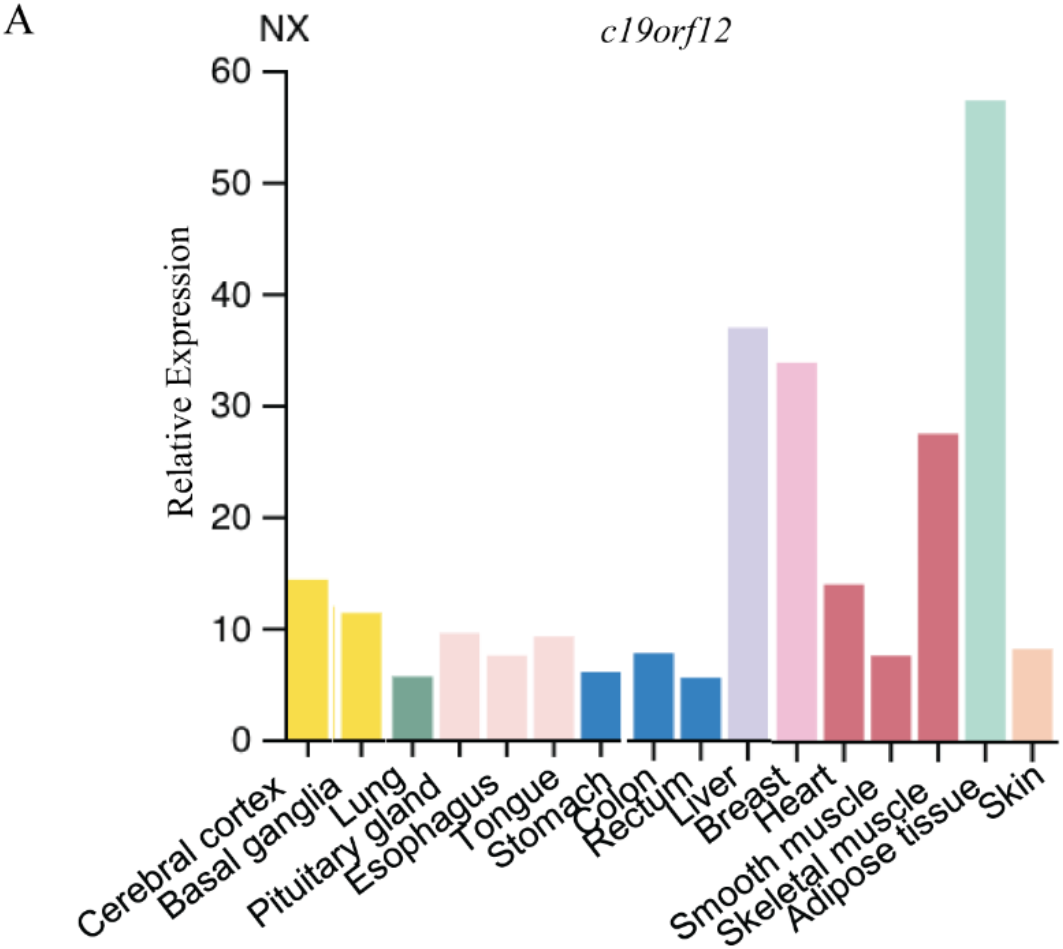
Relative RNA expression in indicated human tissue. (image spliced from RNA expression consensus dataset from human protein atlas).

**SUPPLEMENTAL FILE 1.**
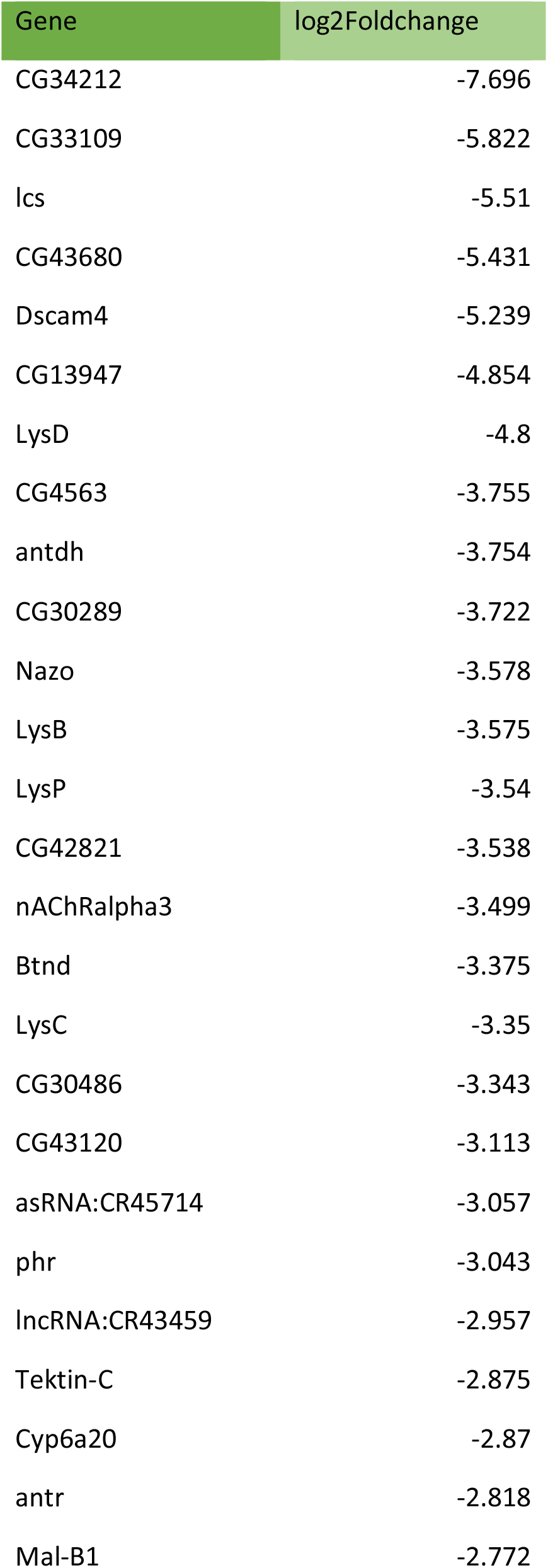
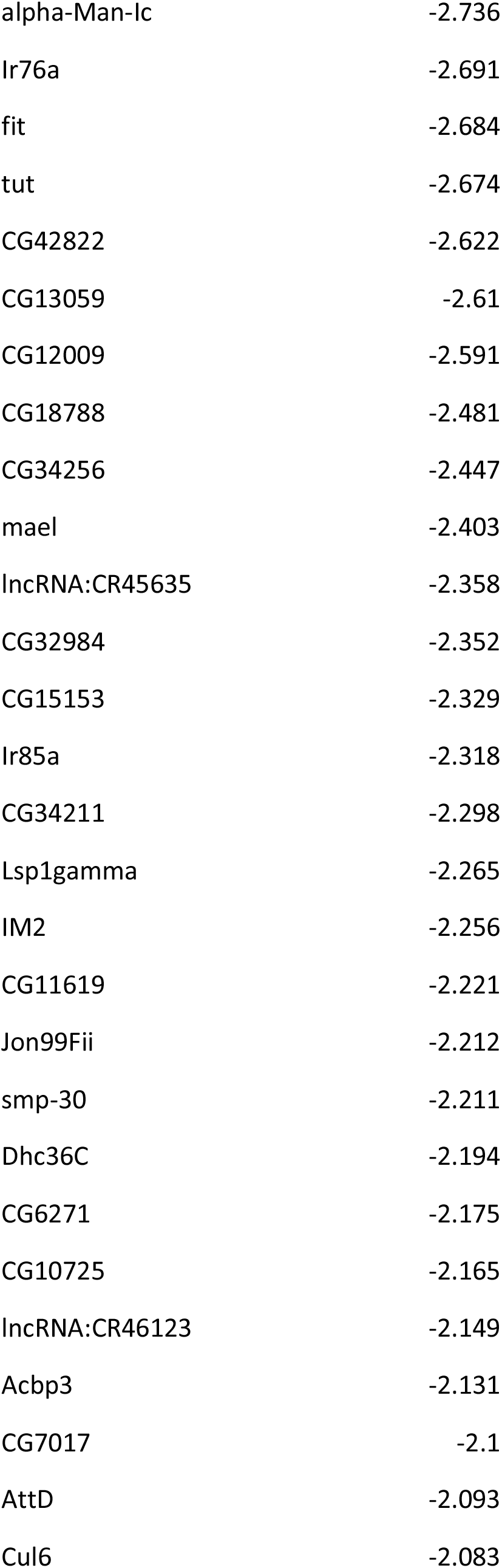
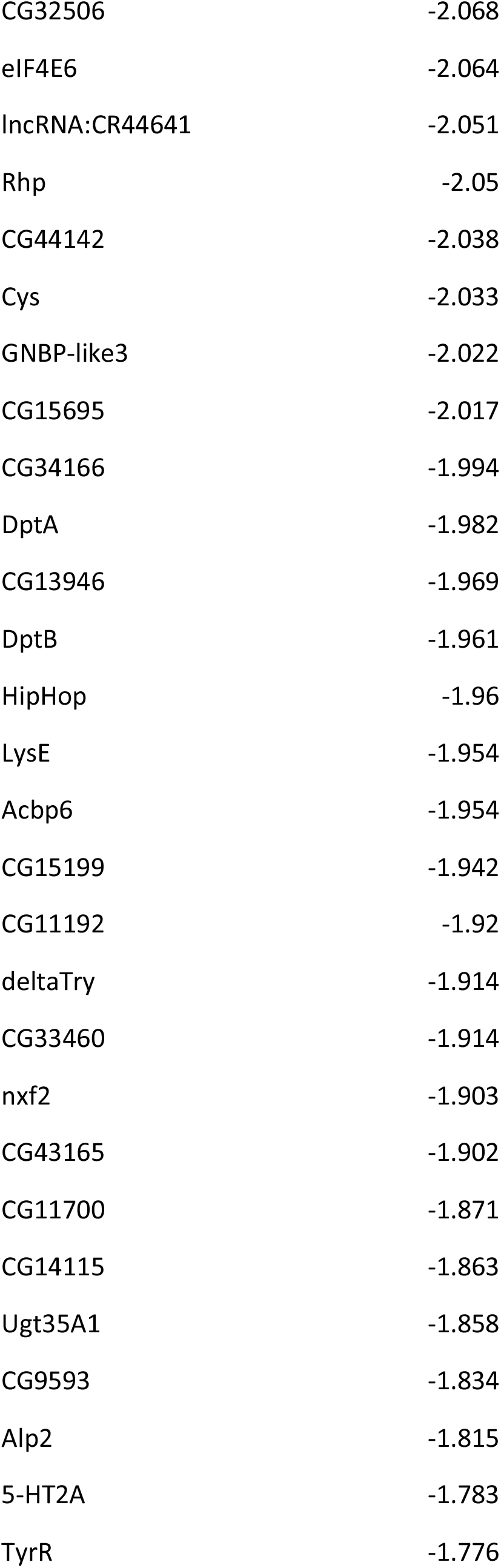
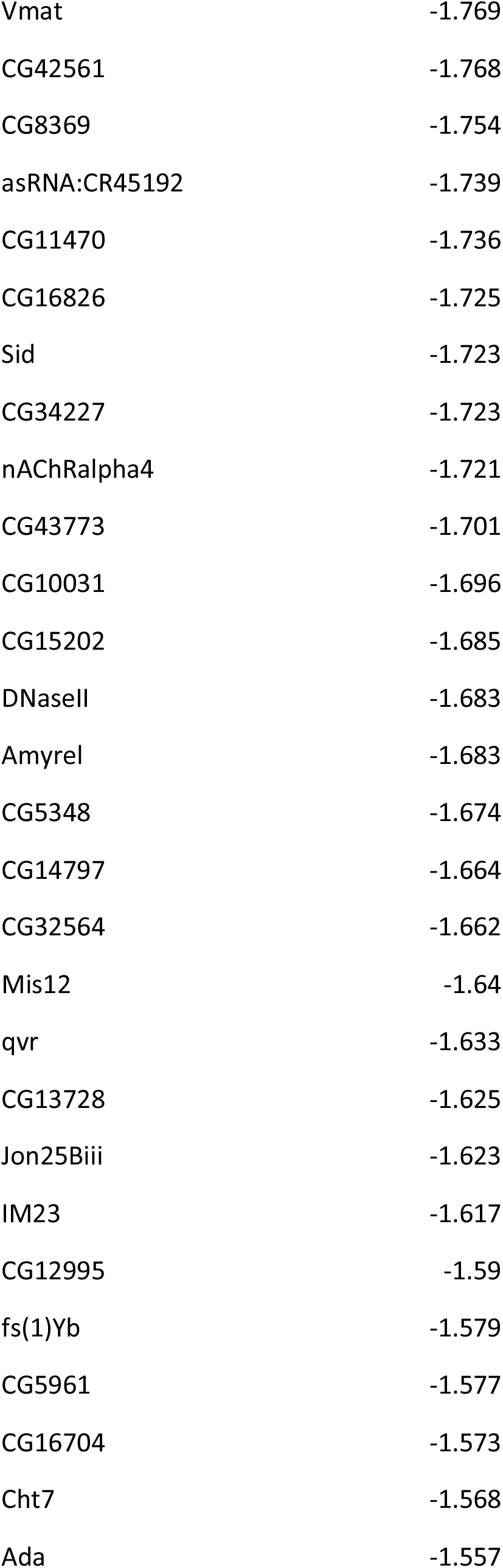
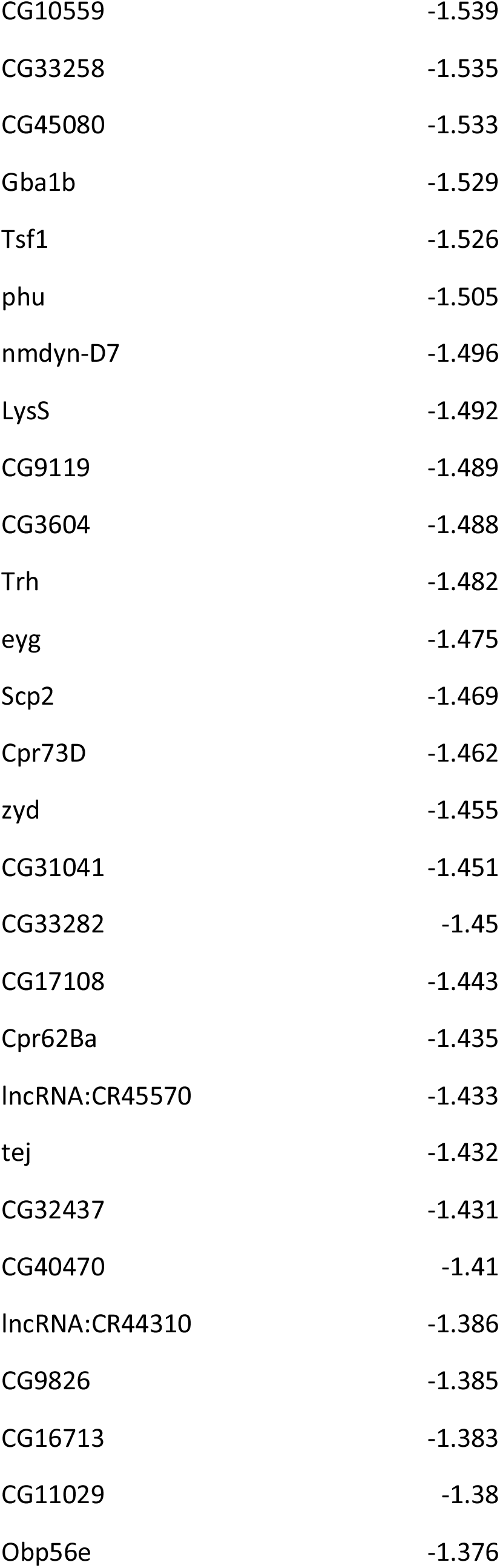
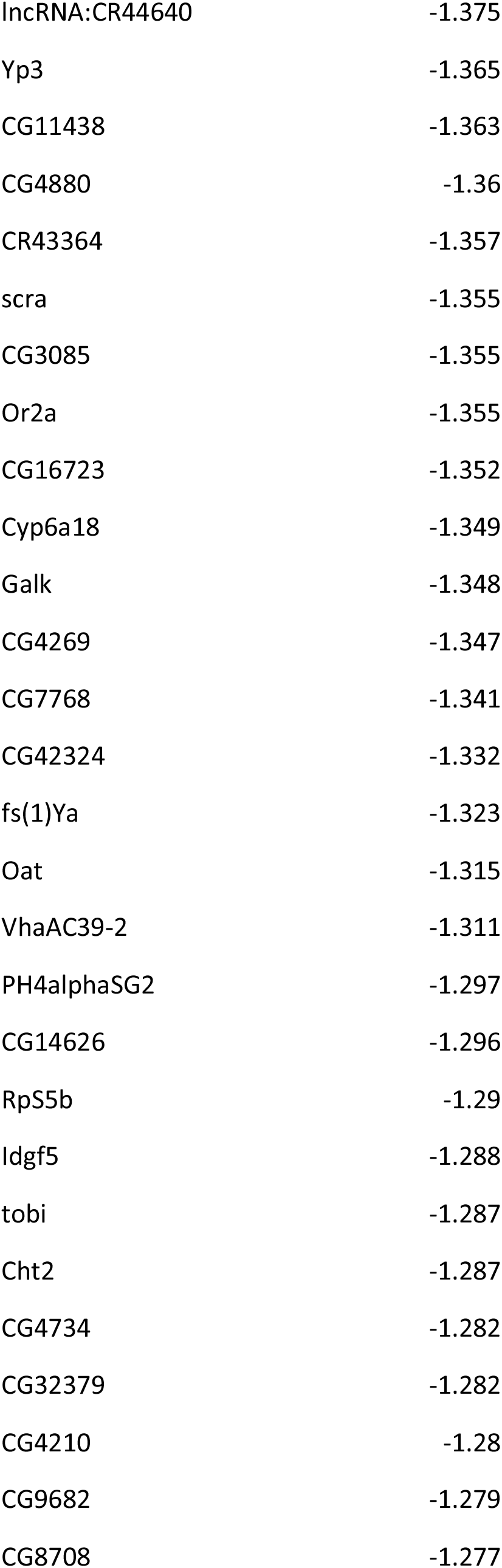
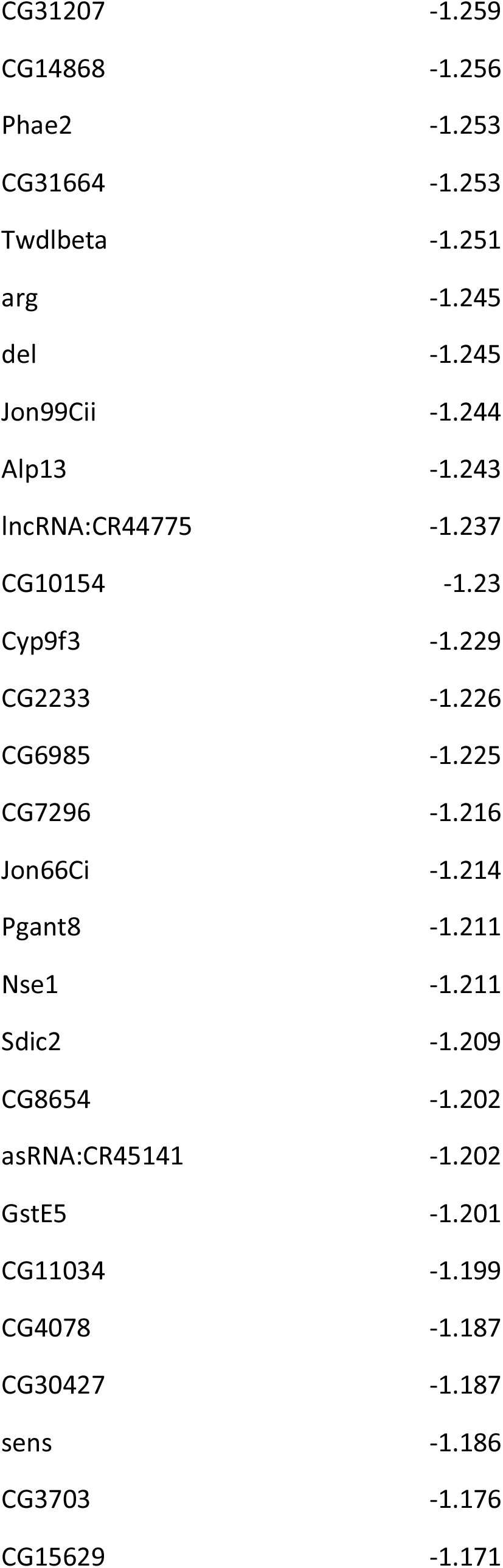
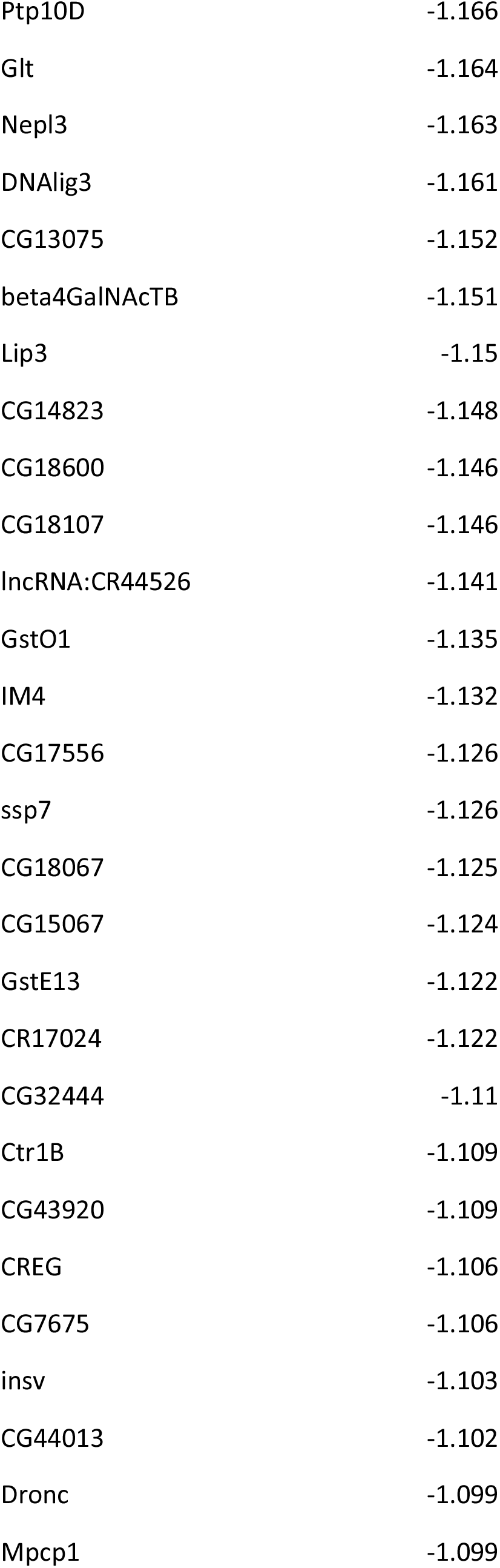
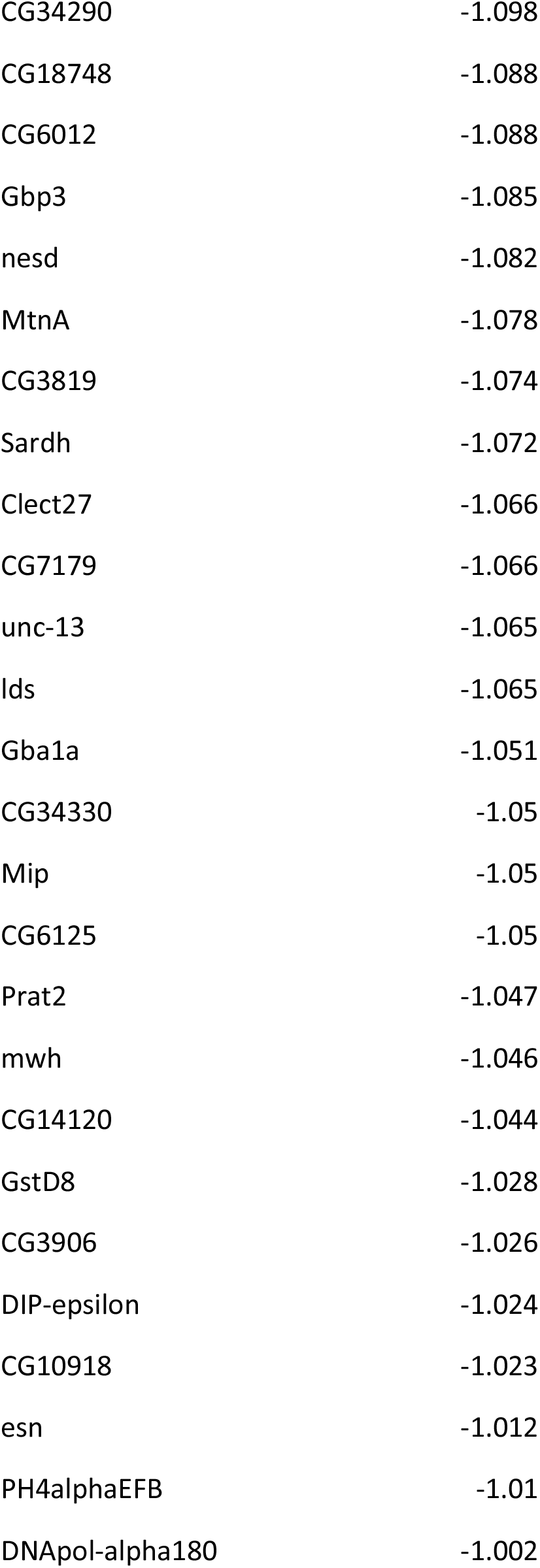
(genes downregulated in *nazo/CG3740* mutants)

**SUPPLEMENTAL FILE 2.** (genes upregulated in *nazo/CG3740* mutants)

| Gene | log2Foldchange |
| --- | --- |
| CG13313 | 5.806 |
| CG11893 | 5.721 |
| asRNA:CR45822 | 5.667 |
| CG32581 | 5.621 |
| asRNA:CR45600 | 5.54 |
| lectin-24A | 5.12 |
| CG4650 | 4.26 |
| CG3397 | 4.117 |
| CG18446 | 4.075 |
| CG43295 | 3.791 |
| Cpr66Cb | 3.759 |
| Hsp70Ba | 3.686 |
| CG7900 | 3.253 |
| l(2)01289 | 3.225 |
| Npc2c | 3.073 |
| CG6912 | 3.006 |
| mthl8 | 2.947 |
| CAH5 | 2.905 |
| Cyp4d1 | 2.898 |
| frm | 2.676 |
| ush | 2.669 |
| lncRNA:CR45138 | 2.655 |
| CG33926 | 2.584 |
| CG6296 | 2.571 |
| CG31741 | 2.54 |
| Arr1 | 2.537 |
| Oseg5 | 2.502 |
| CG31019 | 2.483 |
| CG33346 | 2.475 |
| MtnE | 2.443 |
| tum | 2.331 |
| MtnD | 2.326 |
| CG6738 | 2.306 |
| GILT3 | 2.29 |
| CR46027 | 2.283 |
| CG42399 | 2.268 |
| Jon65Aii | 2.263 |
| lncRNA:CR44138 | 2.253 |
| CG18179 | 2.251 |
| lncRNA:CR43973 | 2.223 |
| CG17570 | 2.171 |
| m | 2.169 |
| CG34138 | 2.127 |
| CG32816 | 2.115 |
| CG30054 | 2.11 |
| ABCA | 2.074 |
| Cpr78Ca | 2.068 |
| Cpr49Ae | 2.05 |
| Spn43Aa | 2.04 |
| CG4830 | 2.038 |
| CG2121 | 2.036 |
| CG42364 | 2.029 |
| CG12744 | 2.015 |
| CG42825 | 2.01 |
| CG12224 | 2.008 |
| CG31464 | 1.956 |
| CG43055 | 1.928 |
| CG31446 | 1.926 |
| CG15431 | 1.901 |
| CG17110 | 1.878 |
| asRNA:CR45044 | 1.846 |
| CG14695 | 1.844 |
| CG31380 | 1.843 |
| CG3635 | 1.836 |
| asRNA:CR45835 | 1.79 |
| CR33013 | 1.776 |
| CG34382 | 1.734 |
| Spn38F | 1.727 |
| CG17752 | 1.686 |
| MFS12 | 1.678 |
| CG18853 | 1.656 |
| Ugt37A1 | 1.656 |
| CG14499 | 1.65 |
| CG3009 | 1.635 |
| rau | 1.619 |
| lncRNA:CR45048 | 1.596 |
| DrsI2 | 1.593 |
| CG4842 | 1.591 |
| CG40298 | 1.565 |
| CG8012 | 1.551 |
| APC7 | 1.536 |
| CG18744 | 1.534 |
| CG4630 | 1.532 |
| CG13676 | 1.53 |
| CG12057 | 1.515 |
| MtnB | 1.499 |
| CG12512 | 1.488 |
| CG13293 | 1.486 |
| CG34291 | 1.477 |
| Cpr50Ca | 1.469 |
| CG33339 | 1.466 |
| Pde9 | 1.456 |
| CR44383 | 1.456 |
| Ca-Ma2d | 1.442 |
| Sfp33A3 | 1.437 |
| CG1139 | 1.434 |
| CG30091 | 1.43 |
| AANATL2 | 1.42 |
| Fst | 1.399 |
| Osi15 | 1.395 |
| Kif3C | 1.394 |
| Ssl2 | 1.384 |
| CG44006 | 1.381 |
| Gld | 1.374 |
| CG43291 | 1.359 |
| CG15254 | 1.357 |
| shd | 1.333 |
| ninaD | 1.327 |
| CG11425 | 1.317 |
| Cyp4g1 | 1.303 |
| Rab26 | 1.299 |
| RYa-R | 1.297 |
| CG43402 | 1.292 |
| D2hgdh | 1.274 |
| CG6277 | 1.268 |
| CG33965 | 1.261 |
| CG11550 | 1.259 |
| hpRNA:CR33940 | 1.256 |
| CG7912 | 1.25 |
| CG11899 | 1.242 |
| Ir25a | 1.242 |
| CG16904 | 1.239 |
| FASN3 | 1.232 |
| CG4080 | 1.231 |
| CG34057 | 1.228 |
| CG7881 | 1.227 |
| l(3)02640 | 1.226 |
| CG17816 | 1.221 |
| CG5550 | 1.218 |
| Lip1 | 1.204 |
| CG9812 | 1.2 |
| CG12194 | 1.189 |
| CG3036 | 1.185 |
| Tina-1 | 1.177 |
| CG10175 | 1.173 |
| CG32641 | 1.171 |
| pst | 1.165 |
| CG14500 | 1.165 |
| CG34215 | 1.164 |
| CG17999 | 1.152 |
| CG6472 | 1.149 |
| Nha2 | 1.147 |
| Acp54A1 | 1.143 |
| Cht5 | 1.136 |
| Amy-d | 1.133 |
| CG12934 | 1.132 |
| CG13067 | 1.128 |
| CG7720 | 1.111 |
| CG30008 | 1.111 |
| Oseg4 | 1.1 |
| IntS9 | 1.098 |
| CG40006 | 1.097 |
| lncRNA:CR43952 | 1.096 |
| rtet | 1.095 |
| lncRNA:CR44024 | 1.092 |
| CG14205 | 1.083 |
| FucTC | 1.077 |
| CG4835 | 1.071 |
| Crys | 1.069 |
| CG11911 | 1.068 |
| CG43367 | 1.062 |
| Ect3 | 1.056 |
| CG31436 | 1.05 |
| CG18586 | 1.049 |
| CG6983 | 1.048 |
| CG8661 | 1.031 |
| CG1698 | 1.024 |
| CG12655 | 1.018 |
| Rgk2 | 1.017 |
| MtnC | 1.012 |
| CG10051 | 1.005 |
| CG10943 | 1.003 |
| CG10383 | 1.002 |
| Ugt302E1 | 1.001 |

